# Novel assays monitoring direct glucocorticoid receptor protein activity exhibit high predictive power for ligand activity on endogenous gene targets

**DOI:** 10.1101/2022.03.29.486227

**Authors:** Laura Van Moortel, Jonathan Thommis, Brecht Maertens, An Staes, Dorien Clarisse, Delphine De Sutter, Claude Libert, Onno C. Meijer, Sven Eyckerman, Kris Gevaert, Karolien De Bosscher

## Abstract

Exogenous glucocorticoids are widely used in the clinic for the treatment of inflammatory disorders and auto-immune diseases. Unfortunately, their use is hampered by many side effects and therapy resistance. Efforts to find more selective glucocorticoid receptor (GR) agonists and modulators (called SEGRAMs) that are able to separate anti-inflammatory effects via gene repression from metabolic effects via gene activation, have been unsuccessful so far. In this study, we characterized a set of functionally diverse GR ligands in A549 cells, first using a panel of luciferase-based reporter gene assays evaluating GR-driven gene activation and gene repression. We expanded this minimal assay set with novel luciferase-based read-outs monitoring GR protein levels, GR dimerization and GR Serine 211 (Ser211) phosphorylation status and compared their outcomes with compound effects on the mRNA levels of known GR target genes in A549 cells and primary hepatocytes. We found that luciferase reporters evaluating GR-driven gene activation and gene repression were not always reliable predictors for effects on endogenous target genes. Remarkably, our novel assay monitoring GR Ser211 phosphorylation levels proved to be the most reliable predictor for compound effects on almost all tested endogenous GR targets, both driven by gene activation and repression. The integration of this novel assay in existing screening platforms running both in academia and industry may therefore boost chances to find novel GR ligands with an actual improved therapeutic benefit.

## Introduction

Synthetic glucocorticoids (GCs) are among the most frequently prescribed drugs worldwide because of their strong anti-inflammatory effects [1]. They are included in the treatment regime for numerous inflammatory diseases, either as topical, inhaled or systemic drugs [2, 3]. Their use is however associated with many side effects, particularly for long-term, high dose systemic treatments [4]. Additionally, many patients develop GC resistance over time [3, 5].

GCs enter cells via passive diffusion and bind the predominantly cytoplasmic glucocorticoid receptor (GR), a nuclear receptor that consists of an intrinsically disordered N-terminal domain (NTD), a DNA-binding domain (DBD), a hinge region and a C-terminal ligand-binding domain (LBD) [6]. Upon ligand binding, GR dissociates from its chaperone complex and migrates to the nucleus. There, it acts as a pleiotropic transcription factor of which the signaling mechanisms are described in detail in [6–8]. Three mechanisms are of particular importance for this study. First, GR forms homodimers which directly bind glucocorticoid response elements (GREs) in target gene promoters or enhancers. This mechanism drives transcriptional activation of anti-inflammatory genes such as glucocorticoid-induced leucine zipper (GILZ) and dual-specificity phosphatase 1 (DUSP1). However, it also contributes to side effects via activation of, among others, tyrosine aminotransferase (TAT) and phosphoenolpyruvate carboxykinase (PEPCK) in the liver, and serum/glucocorticoid-regulated kinase 1 (SGK1) and glutamine synthetase (GS) in other tissues. Second, binding of GR to negative (n)GREs also leads to downregulation of the corresponding target gene. Negative cooperation between the two GR binding sites in an nGRE skews the dynamics towards functionally monomeric GR [9]. The presence of an nGRE in exon 6 of the GR gene itself leads to GC-induced GR downregulation and is one of the mechanisms fortifying therapy resistance [5, 10]. Third, monomeric GR can engage in protein-protein interactions with pro-inflammatory transcription factors such as nuclear factor kappa B (NF-ĸB) and activator protein 1 (AP-1), usually inhibiting their activity and supporting anti-inflammatory effects via a mechanism called tethering. Pro-inflammatory gene suppression was recently found to also occur via direct binding of monomeric GR to cryptic response elements in particular NF-ĸB and AP-1-driven target gene promoters [11, 12]. GR’s actions are influenced further by post-translational modifications (PTMs) and cellular cofactor protein expression and activity profiles [6].

The discrepancy between (mainly desired) monomer-driven tethering and (mainly undesired) dimer-driven effects of GR led to the search for more selective GR ligands (SEGRAMs, SElective GR Agonists and Modulators). SEGRAMs are subdivided into SEDiGRAMs and SEMoGRAMs, aimed at supporting more or less GR homodimer formation than classic GCs, respectively (**Fig. 1**). While SEDiGRAMs could help combat acute inflammation [13], SEMoGRAMs are expected to improve the benefit/risk ratio for treatment of chronic inflammatory disorders because of their limited activity on GRE-driven side effects targets [14].

**Fig. 1.**
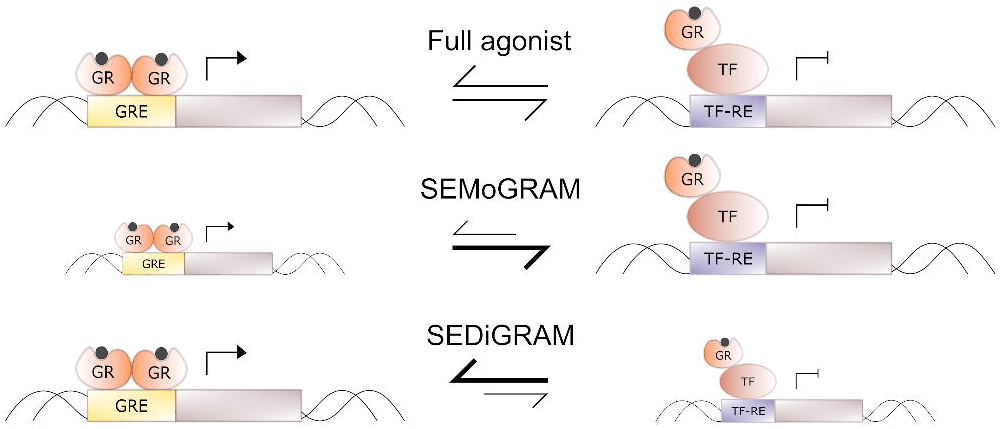
Concept of SEMoGRAM and SEDiGRAM behavior

The search for SEMoGRAMs has proven difficult so far. Although initial studies with compounds such as AL-438, LGD5552, Mapracorat and Fosdagrocorat showed great promise, their pre-clinical (the former two) or clinical development (the latter two) was eventually discontinued (see literature overview in **Supplementary Table 1**). The complexity and context-dependency of GR actions are additional factors complicating successful development of alternatives to classic GCs [15]. Conventional cellular high-throughput screening assays measuring compound activity on reporters driven by GR-responsive promoters potentially oversimplify the complex nature of GR signaling by overlooking gene- and cell-type specificity. Decision-making primarily based on a too limited set of assays should therefore be dissuaded. The need for complementary assays, preferably as cost-effective but with a higher predictive power, thus persists.

The goal of our study was twofold. First, we aimed to investigate to which extent the actions of GR ligands in typical luciferase reporter assays can predict their effects on endogenous GR target genes, in the same and in a different cellular context. Second, we aimed to extend our screening platform (reported in [16, 17]) with novel luciferase-based assays, suitable for medium- to high-throughput screening purposes and, potentially, contributing to a higher overall predictive power. Our findings show that compound effects on the mRNA levels of endogenous GR target genes do not always reflect their actions in screening assays. Surprisingly, we found that quantification of an early event in the signaling pathway of GR, i.e. GR Ser211 phosphorylation, most reliably predicts the behavior of the endogenous GR target genes we studied. We conclude that novel assays focusing directly on the GR protein rather than on its downstream signaling behavior might increase the chances of finding novel SEGRAMs with an actual improved therapeutic benefit.

## Materials & methods

### Compounds, reagents and antibodies

Dexamethasone (Dex, D4902), Cortisol (Hydrocortisone, H4001) and Mometasone Furoate (MF, M4074) were purchased from Sigma-Aldrich. Mapracorat (HY-14864) was purchased from MedChem Express. CompoundA (CpdA) was produced and purchased at the Lab for Medicinal Chemistry (Prof. S. Van Calenbergh, UGhent, Belgium). Prednisolone, Cortivazol, AL-438, LGD5552, RU486, (Fos)dagrocorat, MK-5932, PF-515, AZD2906 and Cpd1-5 were either kind gifts of the respective pharmaceutical companies or re-synthesized based on literature or disclosed patent information [18]. CORT108297 and CORT118335 were a kind gift from CORCEPT Therapeutics. CpdA was stored as a powder at -80 °C. The other compounds were dissolved in DMSO and 10 mM stocks were stored as aliquots at -20 °C. Unless specified otherwise, all compounds were used at a final concentration (f.c.) of 1 µM in 0.1% DMSO, except for CpdA (10 µM) and RU486 (0.1 µM). Recombinant murine (m)TNFα was produced and purified as described earlier [19]. Phorbol 12-myristate 13-acetate (PMA) was purchased from Sigma Aldrich. Mouse anti-GR G-5 (sc393232) and rabbit normal IgG (sc2027) were from Santa-Cruz, and donkey anti-mouse AlexaFluor® 488 and goat anti-rabbit Alexa Fluor 594 were from Invitrogen. Rabbit anti-phosphoGR Ser211 was purchased from Cell Signaling Technology (4161S, lot 2). Mouse anti-Flag M2 and mouse anti-β-actin (AC-15) were from Sigma Aldrich.

### Plasmids

pLV-(IL-6kB)_3_-50hu.IL6P_Luc (in short (NF-ĸB)_3__Luc) and p(GRE)_2_50hu.IL6P-Luc (in short GRE_Luc) have been previously described [20, 21]. pLV-collagenase3_Luc was a kind gift from Dr. E. Canalis (Saint Francis Hospital and Medical Center, Connecticut, USA). pSpCas9(BB)-2A-Puro (px459) was originally created in the Zhang lab (Broad Institute, Cambridge, MA, US) and purchased from Addgene (#62988). NanoBiT plasmids pBiT1.1-N_[TK/SmBiT] and pBiT2.1-N_[TK/LgBiT] were purchased from Promega (cat n° N2014).

### Cell culture

Hek293T cells and all A549-derived cell lines were kept in culture at 37 °C (5% CO_2_), in Dulbecco’s Modified Eagle Medium (DMEM, Invitrogen 41966-021) supplemented with 10% fetal bovine serum (FBS). A549 cells expressing stably integrated (NF-ĸB)_3__Luc, GRE_Luc, collagenase3_Luc or luciferase driven by a constitutive promoter (Const_Luc) were previously described [16]. The A549 (NF-ĸB)_3__Luc GR knock-out (KO) cell line was generated from the A549 (NF-ĸB)_3__Luc cell line via the CRISPR/Cas9 system targeting exon 3 (CATTATGGAGTCTTAA<u>C</u>CTTG) [16]. A549 and cell line variants were, 24 h after cell seeding and 18 h before compound treatment, serum-deprived overnight for all experiments.

### GRE-, ĸBRE- and AP-1-driven luciferase assays

A549 cells with stably integrated luciferase reporters were seeded in black clear-bottom 96-well plates (7,000 cells/well) and treated in triplicate for 6 h with control (0.1% DMSO, 1 µM Dex and 10 µM CpdA) or compound serial dilutions in DMEM without serum. After 1 h, NF-κB-driven reporters were induced with mTNFα (2,000 IU/ml) and AP-1-driven reporters were induced with PMA (100 nM). Cells were lysed in 50 µl 1x Cell Culture Lysis Reagent (CCLR) and luminescence was measured immediately after the addition of 35 µL luciferin substrate. Data were normalized versus 1 µM Dex for each 96-well plate separately, and each assay was conducted at least 3 times to allow robust curve fitting in Graphpad Prism 9.2 (nonlinear fit, 4 parameters).

### Primary hepatocyte isolation

Primary hepatocytes were isolated from male 10-week old C57BL/6 mice by collagenase perfusion, as previously described [22]. 200,000 cells were seeded in collagen-coated 12-well plates. Two hours later, six-hours long compound treatments were started.

### qPCR

A549 and primary hepatocyte cell lysates were homogenized over QIAshredder columns (Qiagen 79656) and RNA was extracted with the RNeasy mini kit (Qiagen 74106) according to the manufacturer’s instructions. 500 ng RNA was converted to cDNA according to the instructions of the cDNA synthesis kit (Biotechrabbit BRO400404). 0.5 µM forward and reverse primers were mixed with LightCycler® 480 SYBR Green I Master Mix (Roche 04 707 516 001) and 25x diluted cDNA for real-time PCR. The primer list is available in **Supplementary Table 2**. Data were analyzed in qBase+. mRNA levels of human target genes in A549 cells were normalized versus hHPRT1, hCyclo and h36B4. mRNA levels of murine target genes in primary hepatocytes were normalized versus mHPRT1 and mB2M.

### NanoBiT

Hek293T cells were seeded in white 96-well plates (10,000 cells/well) and transfected in triplicate 24 h later via calcium phosphate precipitation (50 ng SmBiT-GRα and LgBiT-hGRα, and 10 ng β-actin-gal). 18 h later, the Nano-Glo® Live Cell reagent (Promega) was prepared according to the manufacturer’s instructions. 25 µL was added to each well and the baseline signal was measured using an EnVision 2102 Multilabel reader. Next, the ligands were added and luminescence was measured again after 1 h of treatment. Subsequently, cells were lysed in 20 µL CCLR and 50 µL Galacton-Star™ reagent (ThermoFisher T1056), prepared according to the manufacturer’s instructions, was added. Lysates were incubated for 45-60 min on a shaker at room temperature (RT), and the galactosidase signal was measured on the Envision luminometer. NanoLuc/galactosidase ratios for each separate well were used for further analysis. Data were normalized as fold-difference versus the solvent condition for each biological replicate separately to minimize technical variation.

### Knock-in of Flag-HiBiT at the GR N-terminus

gRNAs targeting the N-terminus of the GR gene (NR3C1, exon 2) were found using the ChOP ChOP search engine and ordered as oligonucleotides (ONTs) from IDT. The ONTs were phosphorylated using T4 kinase (NEB), annealed and cloned into the pSpCas9(BB)-2A-Puro backbone. The single-stranded repair template for Flag-HiBiT knock-in was ordered from IDT as ultramer DNA (sequence in **Supplementary Table 2**). 10^6^ A549 cells were electroporated in duplicate with 5 µg gRNA and Cas9 expression plasmid, and 0.2 nmol ultramer using the Neon™ transfection system from Invitrogen™ (1,000 V, 30ms, 2 pulses). Cells were pooled in a T25 falcon and 24 h later, 1 µg/mL puromycin was added for 48 h. Once the cells reached 90% confluency, single cells were seeded in 96-well plates via manual dilution. Clones were screened for the presence of intact GR and Flag peptide via immunoblotting (described in **Supplementary Information**), and for the presence of the HiBiT tag via HiBiT-based luciferase read-out (described below). The genomic DNA of the positive clones was isolated and the NR3C1 N-terminus was amplified via PCR (35 cycli, 98 °C, 55 °C, 72 °C, primers in **Supplementary Table 1**) and checked by sequencing.

### HiBiT-based quantification of GR levels

This protocol was adapted from [23]. A549 Flag-HiBiT-GR cells were seeded in black 96-well plates (5,000 cells/well) and pre-treated in triplicate with compounds for 1 h before adding 200 IU/mL mTNFα (or DMEM) for 23 h. After these inductions, the medium was removed and replaced by 50 µL DPBS (with Ca^2+^ and Mg^2+^) per well before adding 50 µL NanoGlo® HiBiT lytic buffer supplemented with substrate and LgBiT according to the manufacturer’s instructions (Promega, N3040). The plates were incubated for 10 min on a shaker in the dark before measuring luminescence on the EnVision 2102 Multilabel reader (Perkin Elmer).

### BRET-based quantification of GR Ser211 phosphorylation

This protocol was adapted from [23]. A549 Flag-HiBiT-GR cells were seeded in white 96-well plates (8,000 cells/well) pre-treated in triplicate with compounds for 1 h, before adding 2,000 IU/mL mTNFα for 1 h. Next, the plates were put on ice and washed with 50 µL DPBS (with Ca^2+^ and Mg^2+^) per well. LgBiT (Promega N401A) and rabbit anti-phosphoGR Ser211 were both diluted 1:50 in assay buffer (25 mM Tris-HCl pH 7.8, 150 mM NaCl, 5 mM EDTA, 1% NP-40, HALT protease and phosphatase inhibitors) and 25 µL was added to each well. Assay buffer with 1.48 µg/mL rabbit normal IgG (Santa Cruz sc2027) and 1:50 LgBiT was taken along as negative control. The plates were placed on a shaker platform for 30 min, after which 25 µL assay buffer supplemented with 1:25 furimazine substrate (Promega, N246A) and 1:250 goat anti-rabbit Alexa Fluor 594 was added. Plates were incubated on a shaker platform in the dark for 1 h, after which the NanoLuc and BRET signals were measured on a Tecan infinite M-plex in dual luminescence mode (filter 1: blue1_NB; filter 2: red_NB; integration time 300 ms; settle time 0ms). Each plate was measured in duplicate, after which the individual BRET/NanoLuc ratios were calculated for each well, and the average of the two BRET/NanoLuc ratios were used for further analysis.

### Statistical analysis

All data are expressed as mean ± standard error of the mean (SEM) and are the result of at least three biological replicates. Statistical analysis of all luciferase-based and qPCR data was performed in GraphPad Prism 9.2. Significant differences between compound treatments were evaluated via 1-way or 2-way ANOVA followed by Dunnett’s multiple comparisons test. Heteroscedasticity was evaluated via Brown-Forsythe test, and normality of the residuals was evaluated via Anderson-Darling, D’Agostino-Pearson omnibus, Shapiro-Wilk and Kolmogorov-Smirnov tests. Data were log2-transformed when necessary to obtain a normally distributed, homoscedastic data population. Calculations of Pearson correlations and principal component analysis (PCA) were both conducted in GraphPad Prism 9.2. PCA was performed with parallel analysis, using Monte Carlo simulations on random data (1000 simulations) to filter out principal components capturing technical variation or noise rather than biological variation.

## Results

### Combining GRE-, NF-ĸB- and AP-1-luciferase reporter assay results leads to a refined GR compound classification

For this study, we used a set of 20 GR ligands with maximal functional diversity (**Supplementary Table 1**). All ligands were reported to specifically target GR and the majority were defined as SEGRAMs. Dexamethasone (Dex) was taken along as a reference full agonist, and Compound A (CpdA, non-steroidal) served as an additional anti-inflammatory control compound [21] that targets GR in some cell types but not in others [17, 24, 25]. To confirm that differences in compound activities were not attributed to differences in GR translocation, we evaluated the cellular localization of GR via indirect immunofluorescence in human lung epithelial A549 cells, a well-known GC-responsive cell model. GR was predominantly located in the nucleus for all compounds, except CpdA (**Supplementary Fig. 1**).

Next, we compared the activity of all compounds using three luciferase reporter systems reflecting different GR signaling mechanisms. All reporters were stably integrated in A549 cells and driven by either a GRE (**Fig. 2a-g**), an NF-ĸB response element (ĸB-RE, **Fig. 2h-n**), or an AP-1 response element (TRE, **Fig. 2o-u**). To rule out GR-independent anti-inflammatory effects, all compounds were also tested on an NF-ĸB-driven reporter in GR knocked-out A549 cells (**Supplementary Fig. 2**). No compound showed an effect in the absence of GR, with the exception of CpdA (**Supplementary Fig. 2i**), indicating that the latter can work GR-independently in A549 cells. In parallel, all compounds were tested on a constitutive luciferase reporter to rule out toxicity-mediated effects (**Supplementary Fig. 3**). All data points rendering more than 20% reduction in the activity of this constitutive reporter indicated toxicity and were left out of the concentration-response curves, which were used to calculate the potencies and efficacies of the compounds on all three reporters (**Supplementary Table 3**). Efficacies on ĸB-RE- versus GRE-driven reporters were chosen to classify the compound set into 5 classes: full agonist, partial agonist, antagonist, SEMoGRAM or SEDiGRAM. The criteria for classification are depicted in **Supplementary Fig. 4**. The data on the AP-1-driven Collagenase3_Luc reporter, although largely similar to the ĸB-RE panel results (**Supplementary Table 3**), was excluded from class building since this reporter was strongly downregulated by the known GR antagonist RU486 (**Fig. 2s**). Of note, not all compounds behaved in these assays as expected based on literature (**Supplementary Table 1**).

**Fig. 2.**
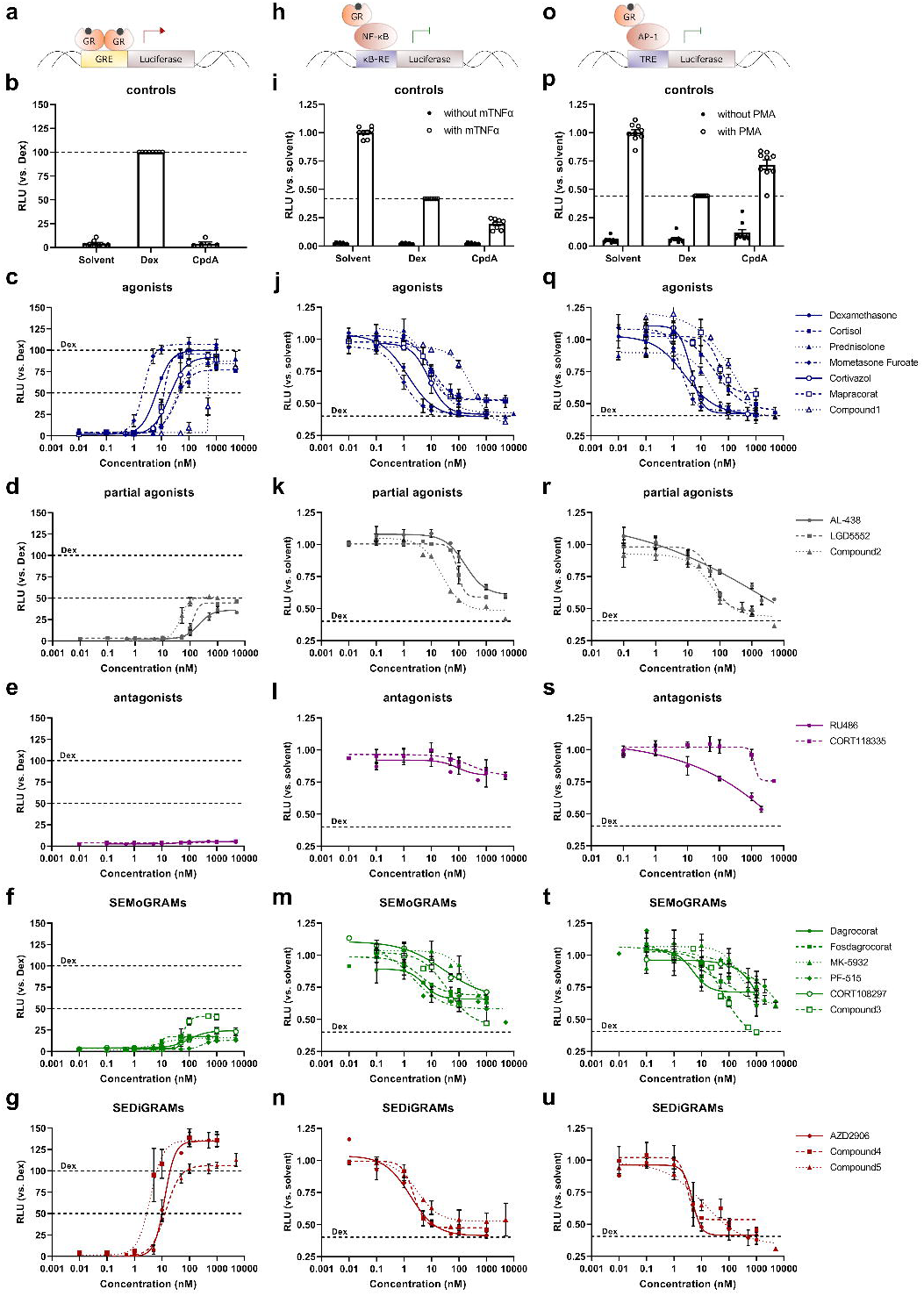
Compound effects on GR-responsive luciferase reporters. A549 cells stably transfected with GRE_Luc (**a-g**), (NF-ĸB)_3__Luc (**h-n**) or Collagenase3_Luc (**o-p**) were seeded in 96-well plates and treated with compound serial dilutions, solvent, 1 µM Dex or 10 µM CpdA for 6h. After 1h, (NF-ĸB)_3__Luc and Collagenase3_Luc were induced with mTNFα (2000 IU/mL) and PMA (100 nM), respectively. All conditions were tested in technical triplicates and data were normalized versus 1 µM Dex for each plate separately. The results from at least 3 biological replicates were used for nonlinear curve fittings in GraphPad Prism 9 (4 parameters). Compounds were classified based on their efficacy on (NF-ĸB)_3__Luc and GRE_Luc (**Supplementary Fig. 3** and **Supplementary Table 3**). RLU, relative luciferase units

### Reporter-based classification does not fully capture SEGRAM behavior on endogenous GR target genes

Next, we assessed whether or not compound classifications based on the limited information conveyed by ĸB-RE- and GRE-driven reporters could predict compound effects on mRNA levels of a panel of endogenous GR target genes driven by either TRE, κB-RE or GRE. We treated A549 cells for 6 h with compounds at high, non-toxic concentrations in the absence or presence of mTNFα. Compound-induced changes in mRNA levels of selected GR target genes largely recapitulated the results and classification from the luciferase assays. Especially the full agonists (blue) and antagonists (magenta) showed very robust and similar profiles on all tested genes from a particular signaling mechanism (**Fig. 3**). The reduced activity of SEMoGRAMs (**Fig. 3a-b**, green) on GRE-driven genes when compared to full agonists was also confirmed. However, strikingly, we failed to confirm a consistently increased activity of SEDiGRAMs (red) over Dex on GRE-driven genes (**Fig. 3a-b**). Although AZD2906 seemed to support higher hFKBP5 and hGS mRNA as compared to Dex, this difference was only statistically significant for hGS in absence of mTNFα (**Supplementary Table 5**). For some compounds, especially LGD5552 (grey), results were highly gene-specific and ranged from antagonist-like behavior on hSGK1 to agonist-like activity on hAngptl4 (**Fig. 3a**).

**Fig. 3.**
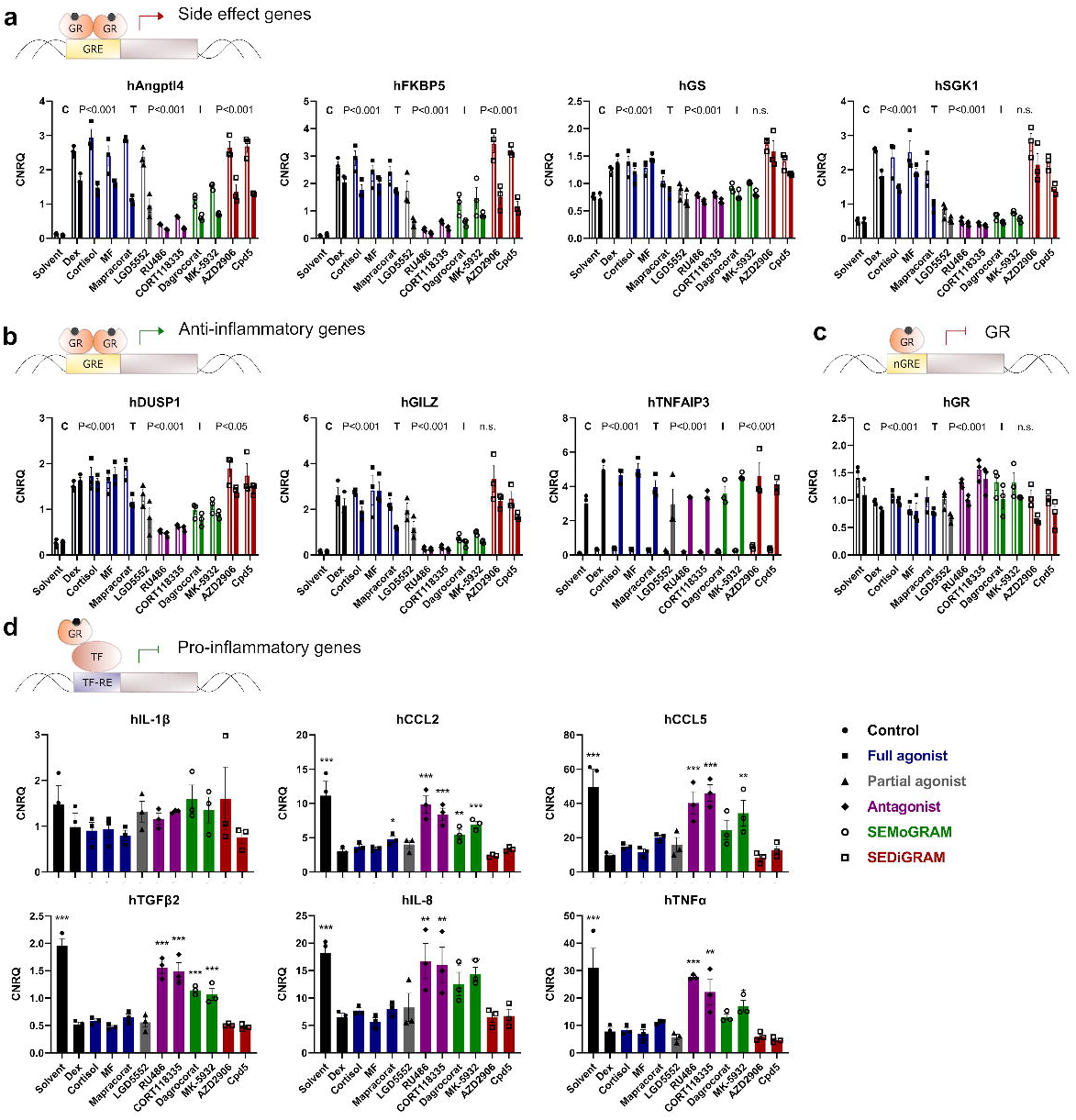
Compound effects on mRNA levels of known GR target genes in A549 cells. A549 cells were pre-incubated with compounds for 1h, before adding mTNFα (2000 IU/mL) for 5h. Total mRNA was isolated, reverse transcribed and used for qPCR analysis. Graphs represent the mRNA levels of the target genes (3 biological replicates), normalized versus hHPRT1, hCyclo and h36B4 mRNA levels. Empty bars indicate conditions without mTNFα, filled bars indicate conditions with mTNFα. **(a)** GRE-driven target genes involved in GC-induced side effects. **(b)** GRE-driven target genes involved in GC-mediated anti-inflammatory effects. **(c)** Downregulation of hGR mRNA via nGRE. **(d)** Indirect GR-mediated downregulation of inflammatory markers. Conditions without mTNFα are depicted in **Supplementary Fig. 5**. (**a-d**) Significant changes versus solvent and Dex were evaluated via 2-way ANOVA followed by Dunnett’s post-hoc testing and are reported in **Supplementary Table 4 and 5. (a-c)** C, compound effect in 2-way ANOVA; T, mTNFα effect in 2-way ANOVA; I, 2-way ANOVA interaction factor. CNRQ, calibrated and normalized relative quantity. (*, P < 0.05; **, P < 0.01; ***, P < 0.001)

mTNFα proved to be a confounding factor for compound activities on GRE-driven target genes, since mTNFα treatment strongly reduced the expression of some GRE-driven genes when compared to conditions without mTNFα (**Fig. 3a-b**). This was reflected in the significant interaction factor between compound and mTNFα treatments in the 2-way ANOVA analysis for hAngptl4, hFKBP5, hDUSP1 and hTNFAIP3. The inhibition of GR signaling by mTNFα, termed reciprocal transrepression [26], has been reported to contribute to side effects and therapy resistance [27]. In contrast to the endogenous target gene setting, this additional effect of mTNFα was not observed with the GRE-driven luciferase reporter (data not shown) and thus seems a feature of more complex promoter constellations.

Since the downregulation of GR mRNA and/or protein levels is an important mechanism contributing to GC resistance, we also determined hGR mRNA levels (**Fig. 3c**). GR mRNA downregulation appeared more prominent with full agonists and SEDiGRAMs compared to antagonists, but the observed differences were rather small and could not support solid conclusions.

Although the SEMoGRAMs scored significantly better than Dex on side effect-related genes (**Fig. 3a**), they could not suppress the mRNA levels of inflammatory markers to the same extent as Dex (**Fig. 3d**). For hCCL5 and hIL-8, SEMoGRAMs combined with mTNFα even failed to cause a significant difference versus mTNFα alone (**Supplementary Table 4**). On the other hand, the partial agonist LGD5552 (grey), was able to suppress most inflammatory markers to the same extent as the full agonists (blue) (**Fig. 3d**) and scored significantly better than Dex on some of the side effect-related genes (**Fig 3a, Supplementary Table 5**). LGD5552 thus yields the best therapeutic index of all tested compounds in A549 cells. Remarkably, SEMoGRAM or SEDiGRAM behavior, as concluded from the reporter gene assays (**Fig. 2**), could not consistently be validated at the mRNA level of relevant endogenous GR target genes.

To unbiasedly study the impact on protein levels following GR activity triggered by the different ligands, we performed mass spectrometry-based shotgun proteomics on A549 cells after 6 h of compound treatment in the presence of mTNFα. Unfortunately, at this time point, we observed no significant changes in protein levels between solvent and the different compounds, not even between solvent and Dex (**Supplementary Fig. 6**). This was most likely due to a lack of sensitivity, since many well-known GR targets could not be identified in this dataset.

### Compound-driven GR activity profiles are only partially conserved between cell types

Since GR actions are known to be highly gene- and tissue-specific [28, 29], we next evaluated compound effects in murine primary hepatocytes as proxies for cellular metabolic side effect mechanisms. While the compound activity profiles were very similar as in A549 cells for some genes (e.g. mGR, mAngptl4 and mSGK1), other genes showed very strong differences (e.g. mGS, mDUSP1) (**Fig. 4**). The latter underwent only minor regulation in primary hepatocytes after 6h of compound treatment, in contrast to 6 h treatments in A549 cells (**Fig. 3a-b**). Another contrasting finding compared to A549 cells is that the as a SEDiGRAM classified Cpd5 (red) induced significantly higher mRNA levels of mPEPCK and mAngptl4 when compared to Dex (**Fig. 4a**). However, mPEPCK is most likely not a reliable parameter for a refined classification of GR ligands, since the effects of Dex and RU486 on this gene could not be distinguished (**Fig. 4a)**. Strikingly, in primary hepatocytes, Mapracorat (blue) rather behaved as a partial agonist or even as a SEMoGRAM, and not as a full agonist as in A549 cells (**Fig. 2, Fig. 3**). Mapracorat’s partial agonist behavior was most pronounced on mFKBP5, mSGK1, mFam107a and mTAT (**Fig. 4a-b**). LGD5552 (grey) supported even lower mRNA levels of FKBP5 and Angptl4 in primary hepatocytes than in A549 cells (**Fig. 3a, Fig. 4a**). These data clearly point to the importance of testing novel SEGRAMs on a representative panel of relevant target genes in relevant target cells.

**Fig. 4.**
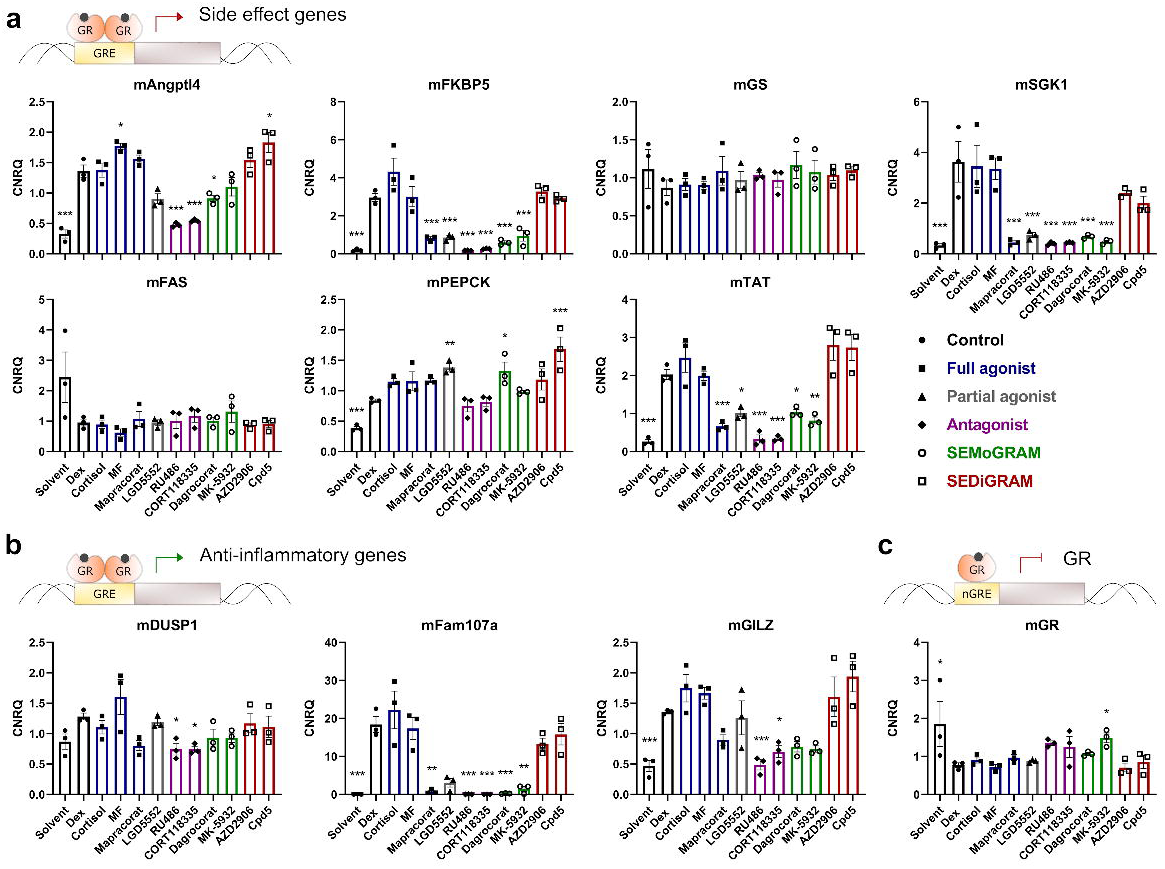
Compound effects on mRNA levels of known GR target genes in primary hepatocytes. Primary hepatocytes were isolated from male 10-week old C57BL/6 mice by collagenase perfusion, allowed to attach for 2h and subsequently treated with compounds for 6h. Total mRNA was isolated, reverse transcribed and used for qPCR analysis. Graphs represent the mRNA levels of target genes (3 biological replicates), normalized versus mHPRT1 and mB2M mRNA levels. Statistically significant differences versus Dex were evaluated via one-way ANOVA followed by Dunnett’s post-hoc testing on log2-transformed data (*, P < 0.05; **, P < 0.01; ***, P < 0.001). **(a)** GRE-driven target genes involved in GC-induced side effects. **(b)** GRE-driven target genes involved in GC-mediated anti-inflammatory effects. **(c)** Downregulation of mGR mRNA via nGRE. CNRQ, calibrated and normalized relative quantity

### Novel luciferase-based assays allow evaluation of compound effects directly on the GR protein

Partial discrepancies between compound activities on GR-driven luciferase reporters versus endogenous target genes emphasize a need for novel screening assays, preferentially with higher predictive power than currently existing assays while still amenable to high-throughput screening. For instance, we observed limited discriminatory power when validating compound classification based on the GRE-driven luciferase reporter. Compounds classified as SEDiGRAMs did not consistently recapitulate SEDiGRAM behavior on GRE-driven mRNAs in A549 cells or primary hepatocytes, highlighting the need to reinforce the current screening platform. As a next step, we reasoned that reinforcement of the current screening platform may improve on predictive power.

As the entire concept of SEMoGRAMs and SEDiGRAMs is based on their differential capacity to induce GR dimerization, we implemented the NanoBiT technology to quantify GR dimerization. Two constructs were cloned, linking the two different portions of a split NanoLuc luciferase (SmBiT and LgBiT) to the GR N-terminus. Given the low affinity of SmBiT and LgBiT for one another, reconstitution of functional NanoLuc in intact cells relies entirely on GR dimerization. First tests using a GR knock-out A549 cell line were hampered by poor transfection efficiencies. Therefore, we switched to the easily transfectable Hek293T cells, which express negligible levels of functional endogenous GR. NanoBiT assay results revealed that none of the tested SEDiGRAMs (red) induced higher GR dimerization levels than Dex or the other classic agonists (blue) (**Fig. 5a**), which is in line with most of the qPCR data. Encouragingly, both antagonists (magenta) and the majority of the SEMoGRAMs (green) behaved entirely as expected and caused significantly less GR dimerization than Dex, hereby also confirming the validity of this assay. The dimerization capacity of GR when triggered by compounds from these two groups was highly alike, which is in line with the hypothesis from Hu and colleagues that true SEMoGRAM compounds may rather exhibit a GR antagonist-like profile [30]. Partial agonists displayed intermediate, compound-specific, effects on GR dimerization.

**Fig. 5.**
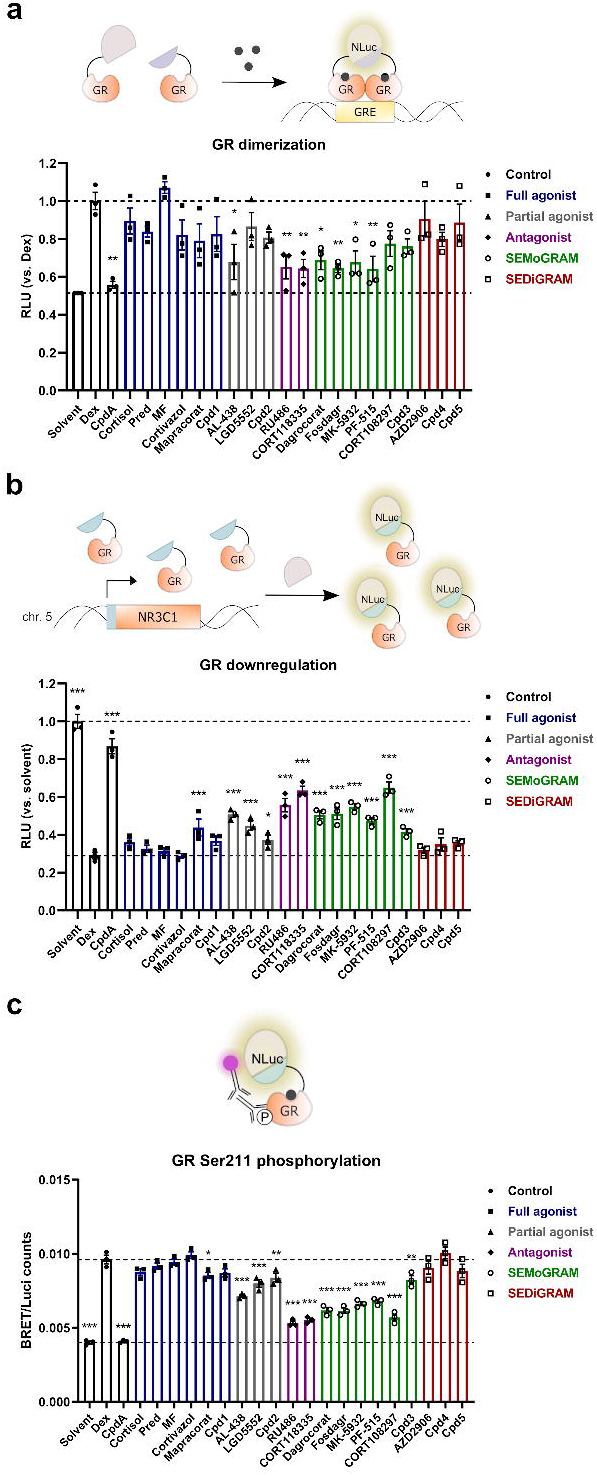
Novel screening assays directly evaluating compound effects on GR protein **(a)** Quantification of GR dimerization in Hek293T cells using the NanoBiT system. Hek293T cells were transfected with SmBiT-GRα, LgBiT-GRα and β-actin_gal. NanoLuc and galactosidase activity was measured 24h later, after 1h compound treatment. NanoLuc/Galactosidase ratios were normalized versus solvent for each plate individually. The graph represents relative dimerization versus Dex. **(b)** Quantification of GR levels in an A549 Flag-HiBiT-GR knock-in cell line. A549 cells were engineered using CRISPR-Cas9 and a single-stranded DNA repair template to introduce a dual Flag-HiBiT tag at the GR N-terminus. A549 Flag-HiBiT-GR cells were treated with compounds for 24h and NanoLuc activity was measured using the NanoGlo® HiBiT lytic detection system. Data are expressed as fold versus solvent. **(c)** Quantification of GR Ser211 phosphorylation levels via BRET. A549 Flag-HiBiT-GR cells were pre-incubated with compounds for 2h. Lysates were incubated with anti-GR and LgBiT for 30 min, followed by addition of furimazine substrate and anti-rabbit Alexa Fluor 594 for 60 min. BRET/NanoLuc ratios were calculated for each well separately and used for further analysis. Data reporting compound effects in presence of mTNFα are available in **Supplementary Fig. 8**. (**a-c**) All conditions were measured in technical triplicates. Statistically significant changes versus Dex were evaluated via ordinary one-way ANOVA (**a**-**b**) or 2-way ANOVA (**c**) with Dunnett’s post-hoc testing on log2-transformed data (3 biological replicates) (* P < 0.05; ** P < 0.01; *** P < 0.001). (**c**) Results from post-hoc testing are provided in **Supplementary Table 6**. RLU, relative luciferase units

Next to therapy-associated side effects, acquired GC resistance due to gradually declining GR protein levels also overshadows the benefit of GC therapy. Since the monitoring of GR downregulation via qPCR (**Fig. 3c, Fig. 4c**) or reporters (data not shown) lacked sensitivity and discriminatory power, we developed an alternative assay for fast and quantitative monitoring of GR protein levels. We used a CRISPR Cas9-based knock-in system to introduce dual Flag and HiBiT tags at the N-terminus of endogenous GR in A549 cells. The HiBiT tag is a small portion of NanoLuc with very high affinity for LgBiT, ensuring reconstitution of functional NanoLuc upon addition of recombinant LgBiT to cellular lysates. The resulting A549 Flag-HiBiT-GR (FH-GR) cell line nicely recapitulated Dex-mediated GR downregulation while control experiments confirmed functionality of the tagged GR (**Supplementary Fig. 7**). A549 FH-GR cells were treated with GR ligands for 24 h in the presence or absence of mTNFα. Since mTNFα did not have any significant effect on the regulation of GR levels, we only used the data obtained without mTNFα added. **Fig. 5b** shows that the HiBiT-based read-out is a highly robust and sensitive approach to monitor GR protein levels, which were the lowest after treatment with full agonists (blue) and SEDiGRAMs (red), in line with GR mRNA levels (**Fig. 3c, 4c**). Mirroring qPCR and NanoBiT data, effects of SEDiGRAMs and full agonists were indistinguishable. Both antagonists (magenta) and SEMoGRAMs (green) downregulated GR to a lesser extent than Dex, which indicates that SEMoGRAMs not only have the potential to reduce side effect burdens, but could also be better than conventional GCs in the context of therapy resistance. Notably, all compounds, even the antagonists (magenta), still caused significant GR downregulation compared to solvent, except CpdA.

HiBiT tagging also allows quantification of PTMs of the protein of interest via bioluminescence resonance energy transfer (BRET) analysis [23]. We opted for GR Ser211 phosphorylation, as an important hallmark of GRE-driven activity [31]. All compounds (except CpdA, as previously reported [17, 21]) significantly increased the levels of phosphorylated Ser211 compared to the solvent condition, both in absence or presence of mTNFα (**Fig. 5c, Supplementary Fig. 8**). In line with the NanoBiT results (**Fig. 5a**), the compounds largely clustered together in two large groups, with the full agonists (blue) and the SEDiGRAMs (red) causing the highest levels of Ser211 phosphorylation, and the antagonists (magenta) and the SEMOGRAMs (green) leading to lower levels. Partial agonists again fluctuated between these groups (**Fig. 5c, Supplementary Fig. 8, Supplementary Table 6**). In contrast to the downregulation of GR protein levels, mTNFα showed a small, yet statistically significant reduction of the Ser211 phosphorylation levels, in line with the reported reciprocal inhibition between TNF and GR pathways [31]. SEDiGRAMs showed no statistically significant differences compared to Dex, reflecting their effects on mRNA levels of most GRE-driven target genes (**Fig. 3a-b, Fig. 4a-b**), but contradicting GRE-driven luciferase reporter data used to initially classify them as SEDiGRAMs (**Fig. 2a-g**). Mapracorat caused significantly less Ser211 phosphorylation than Dex and clustered together with the full agonists, in line with other A549 data but in contrast to its effects in primary hepatocytes.

### GR Ser211 phosphorylation levels correlate strongest with outcomes on endogenous GR target genes

Since one of the main goals of this research was to evaluate how well actions of compounds on one screening parameter could predict the outcome on another parameter, we calculated pairwise Pearson correlations for all luciferase-based assays and the qPCR data using a compound subset that represents each class (**Fig. 6**). First, we compared how well each of the compound activities on the luciferase-based assays correlated with the effects on the mRNA levels of GRE-driven target genes (**Fig. 6a)**. Overall, correlation coefficients were high. The major exceptions were the mRNA levels of mGS, mDUSP1 and mPEPCK, which responded poorly to the compound treatments (**Fig. 4a-b**). The effects on essentially all GRE-driven qPCR targets correlated best with the BRET assay data (**Fig. 6a**, magenta box) quantifying GR Ser211 phosphorylation levels, more than with the GRE-driven luciferase reporter data. The only exceptions were hGS and hSGK1 mRNA levels, which correlated slightly better with the GRE-driven luciferase reporter read-out (**Fig. 6a**).

**Fig. 6.**
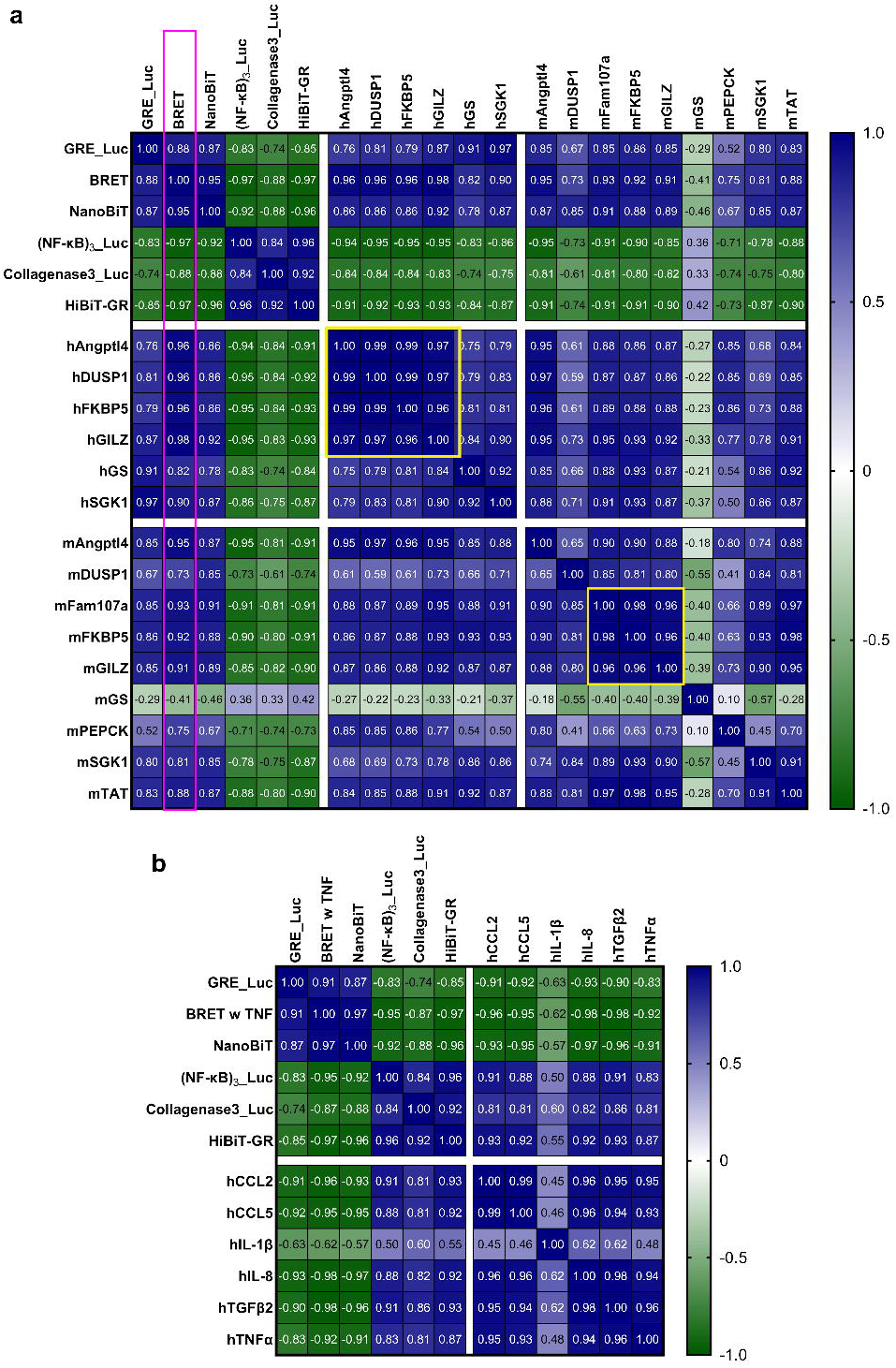
Correlation studies of the data reported in **Fig. 2-5**. (**a**) Heath map representing the Pearson correlations between the luciferase-based assays and all tested genes representing GRE-driven gene activation. We included the BRET data in absence of mTNFα (**b**) Heath map representing the Pearson correlations between the reporter gene assays and all tested AP-1- and/or NF-ĸB-driven inflammatory genes. We included the BRET data in presence of mTNFα (**a, b**) Pearson correlations were calculated based on the mean values of the log2-transformed data in GraphPad Prism 9. Magenta box highlights correlation coefficients between BRET and other assays; cyan boxes highlight qPCR targets with highest correlation coefficients

Similarly, GRE-driven target mRNA levels also correlated better with NanoBiT dimerization results than with the GRE-driven reporter data. However, the NanoBiT assay did not score as well as the BRET assay, indicating that GRE-driven transcriptional activity is linked more closely with Ser211 phosphorylation than with GR dimerization. Most strikingly and quite unexpectedly, both the ĸB-RE-driven reporter and the HiBiT-based GR-downregulation assay had a higher predictive power for the outcomes on the mRNA levels of GRE-driven genes than the NanoBiT and GRE-driven reporter assay. This was reflected in high mutual (absolute) correlation coefficients for the outcomes of the ĸB-RE-driven reporter assays, HiBiT assays, BRET and NanoBiT (**Fig. 6a**). This was still observed when comparing their effects on the larger dataset with 20 compounds (**Supplementary Fig. 9a**).

Along the same lines, high Pearson correlations were observed between the compound effects in BRET and NanoBiT assays on the one hand, and the mRNA levels of pro-inflammatory markers on the other (**Fig. 6b**). The former two assays actually correlated better with the effects on the mRNA levels of pro-inflammatory markers than the ĸB-RE- and AP-1-driven luciferase reporters, even though GR dimerization and Ser211 phosphorylation are essential for GRE-driven transcriptional activity and not typically linked to tethering mechanisms [31–33]. In support, very high absolute correlation coefficients were noted between some endogenous anti-inflammatory and pro-inflammatory genes, such as hGILZ and hIL-8 (|r| = 0.98, **Supplementary Fig. 9b**).

Another interesting observation was that the mRNA levels of some endogenous target genes showed markedly similar compound-induced expression profiles, as verified by very high Pearson correlations (r ≥ 0.95). As such, effects on hAngptl4, hDUSP1, hFKBP5 and hGILZ clustered in the A549 data, while mFam107a, mFKBP5 and mGILZ clustered in the data obtained from murine primary hepatocytes (**Fig. 6a**, cyan boxes). Additionally, compound effects on hGILZ correlated strongly with those on mAngptl4 and mFam107a (**Fig. 6a**).

Given the discrepancies between data from classic luciferase reporters (**Fig. 2**) and all other assay data, we wondered how the compounds would cluster when including all the data, and whether we would still be able to distinguish the five compound classes. Therefore, we performed a Principal Component Analysis (PCA) on the dataset consisting of 20 compounds that were screened in the luciferase-based assays (**Fig. 7a**). A second dataset consisted of 10 compounds and their effects on all luciferase reporters and mRNA levels of endogenous target genes (**Fig. 7b**). PCA on both datasets revealed that one principal component (PC1) sufficed to capture all biological effects, explaining 88,9% of variation in the first dataset (**Fig. 7a**) and 84,5% in the second (**Fig. 7b**). Comparisons with random datasets using Monte Carlo simulations revealed that PC2 represented noise instead of biologically relevant effects, leaving PC2 out of the final graphs. The luciferase-based data alone proved insufficient to distinguish the different compound classes and largely separated them in only two groups (**Fig. 7a**), one containing the antagonists and SEMoGRAMs, and another the full agonists and SEDiGRAMs, as also observed in the NanoBiT and BRET profiles (**Fig. 5a, c, Supplementary Fig. 8**). When including the qPCR data in the PCA, four groups could be distinguished: antagonists, SEMoGRAMs, partial agonists and full agonists/SEDiGRAMs (**Fig. 7b**). The latter two were inseparable from each other. Even though the two SEMoGRAMs (Dagrocorat and MK-5932) nicely clustered, their general activity profile appeared to be intermediate between antagonists and partial agonists without causing the shift that would be expected when affecting the stoichiometry of GR dimerization (**Fig. 7c**). This theoretical shift was not only absent for SEMoGRAMs but also for SEDiGRAMs, as exemplified in the correlation plot between regulation of hIL-8 mRNA levels (representing monomer-driven effects) and hGILZ mRNA levels (representing dimer-driven effects (**Fig. 7c**), Based on all data in the model systems used here, Mapracorat rather classified as a partial agonist than a full agonist.

**Fig. 7.**
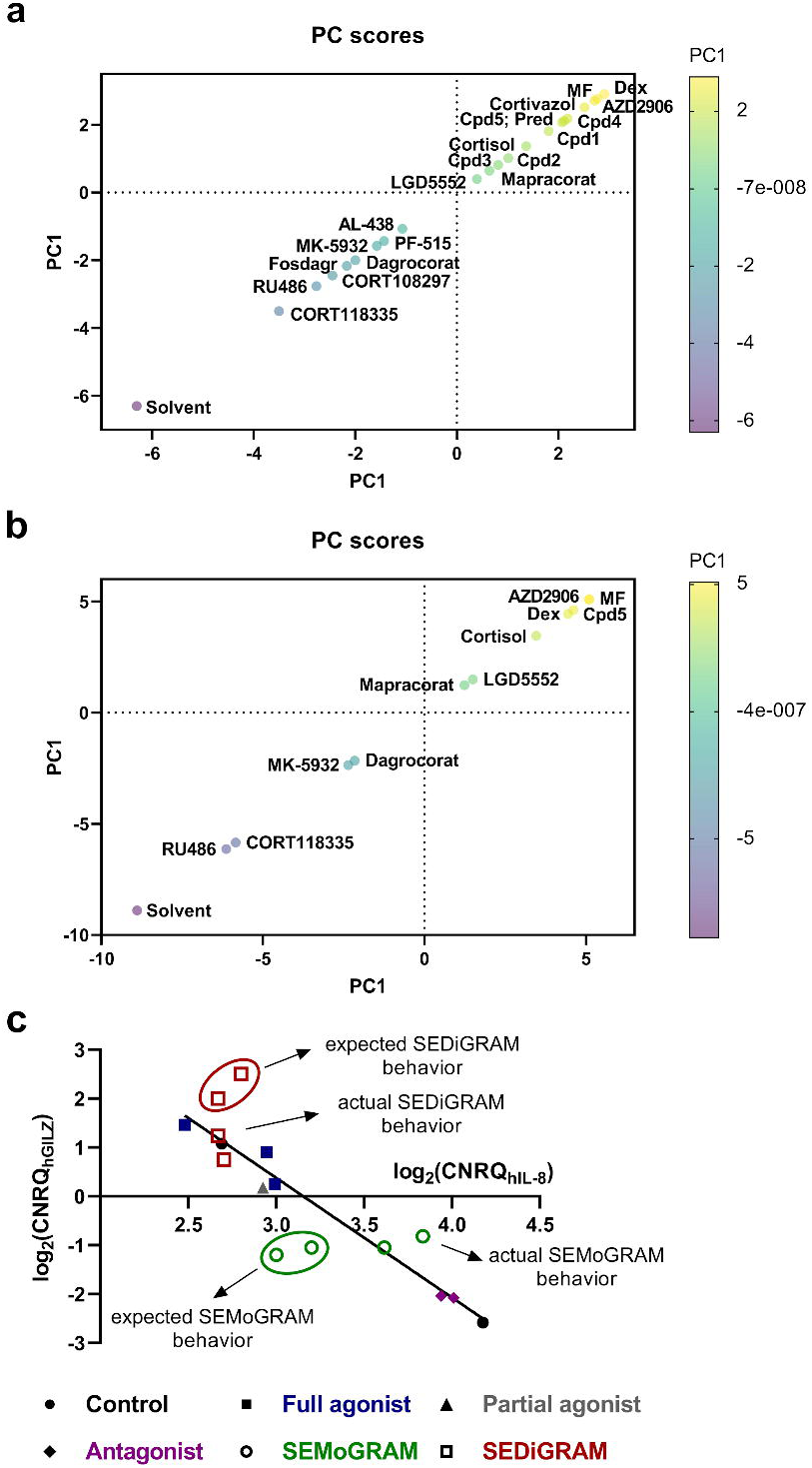
Expected versus actual SEGRAM behavior **(a-b)** Principal Component Analysis (PCA) of the data reported in **Fig. 2-5 (a)** PCA based on the luciferase data with 20 compounds. **(b)** PCA based on the luciferase and qPCR data with 10 compounds **(c)** expected versus actual behavior of SEMoGRAMs and SEDiGRAMs, exemplified on mRNA levels of hGILZ and hIL-8

## Discussion

In this study, we closely examined currently available assays for their ability to reliably predict ligand behavior. Even though a minimal set of reductionist-type assays may be cost-effective, it is often at the expense of discriminatory capacity. It also complicates decision-making processes of what hit compounds should be taken forward to lead development programs. Novel, sensitive cell-based assays that evaluate distinct properties of the GR signaling pathway could solve this lack of discriminatory power. Overall, we developed an advanced GR assay platform to allow a better-informed prioritization of compounds following first-stage high-throughput screenings. Our findings may boost active research on selective GR ligand development.

To substantiate the unmet need for improved GR screening systems, we characterized a set of functionally diverse GR ligands (**Supplementary Table 1**), first using a panel of commonly used luciferase-based reporter gene assays solely capturing GR-driven gene activation and gene suppression events (**Fig. 2**). We expanded this minimal set with novel read-outs quantifying GR protein levels, dimerization and phosphorylation status (**Fig. 5)**. To minimize bias due to cell- and tissue-specificity and to ensure maximal comparison across all assays, we optimized all assays in the same cellular system, A549 cells, except for NanoBiT, which we could only get to work in Hek293T cells. We validated the outcomes by monitoring the mRNA levels of a subset of known GR target genes in A549 cells and primary hepatocytes (**Fig. 3, Fig. 4**). To our knowledge, this is the first study comparing the cellular behavior of more than ten published SEGRAMs in a uniform cellular context on different GR signaling parameters, in parallel. Our analyses support that the final screening platform combined with mRNA validations effectively result in a more refined GR compound classification system (**Fig. 7**).

Throughout the study, surprising observations came to light when comparing compound behavior across assays. First, we discovered that a TRE-driven luciferase reporter (AP-1 mediated) may be an unreliable parameter for compound classification, since it was strongly downregulated by the known GR antagonist RU486 (**Fig. 2s**). We also found that classification of compounds solely based on GRE-driven and ĸB-RE-driven luciferase reporter data (**Fig. 2, Supplementary Fig. 4**) did not always align well with mRNA levels of various representative GR target genes supposedly regulated via similar mechanisms (**Fig. 3, Fig. 4**). For instance, the increased GRE-driven activity of compounds classified as SEDiGRAMs could not be validated on any of the tested endogenous GR target genes in A549 cells (**Fig. 3**). We did observe SEDiGRAM activity of Cpd5 on the mRNA levels of mAngptl4 and mPEPCK in primary hepatocytes (**Fig. 4a**), but the latter gene could not distinguish agonist from antagonist effects and is therefore not a reliable screening parameter. This was also reflected in the poor correlation between the effects on mPEPCK and other GRE-driven targets (**Fig. 6a**). Given the well-documented context-dependency of GR actions inferring a role for the chromatin landscape [6, 34], the divergences between the reporter data and endogenous target gene profiles were not completely unexpected [15]. Neither were the observed discrepancies between our compound classifications and the reported compound behaviors as published in literature (summarized in **Supplementary Table 3**). Both demonstrate the importance of validating the results of high-throughput screening efforts on endogenous target genes in relevant cell models.

When comparing the compound effects on all tested target genes in A549 cells and primary hepatocytes, we observed similarities as well as differences. The compound response profile on the mRNA levels of hGILZ clustered together with the effects on hAngptl4, hDUSP1 and hFKBP5 in A549 cells, as well as with mAngptl4 and mFam107a in murine primary hepatocytes (**Fig. 6a, Supplementary Fig. 9b**). This indicates that the mRNA expression profile of hGILZ could be used as a predictor for the effects on these five other genes, at least within these two cellular models. However, other target genes showed strong regulatory differences between the two cell types. For GS and DUSP1, this effect was probably due to general differences in expression levels (**Fig. 3a-b, Fig. 4a-b**). For other genes, the cell type-specific differences were limited to one particular compound, for instance the differential effects of Mapracorat on FKBP5, GILZ and SGK1 between the two cell types (**Fig. 2a, Fig. 3a**). Mapracorat generally behaved as a full agonist in A549 cells (**Fig. 2, Fig. 3, Fig. 5**), while it rather acted as a partial agonist or even a SEMoGRAM in primary hepatocytes (**Fig. 4**). The latter observation is in line with earlier studies showing that subcutaneously injected Mapracorat caused significantly less activation of mTAT *in vivo* when compared to mometasone furoate (MF) [35]. Mapracorat also acted as an improved ligand in rabbit models of dry eye disease, where it caused significantly less increase in ocular pressure than Dex [36]. The cell-specific actions of Mapracorat demonstrate the need to screen in a relevant cellular context. Another interesting example showing context-dependent actions of GR ligands, is CORT118335. This compound was shown to act as a partial GR agonist *in vivo* in a mouse NAFLD model [37], yet consistently behaved as an antagonist in our assays in both cell models.

When comparing the overall correlation coefficients between the mRNA levels of all evaluated target genes in A549 cells (**Fig. 6b, Supplementary Fig. 9b**), we observed that mutual correlations between effects on anti-inflammatory targets (subject to indirect downregulation) were even higher than between GRE-driven target genes. This could indicate that the latter are more prone to gene- and tissue-specificity than the former [38]. This also implies that GRE-driven compound activities could be more difficult to predict based on a limited screening panel. While tethering-based actions of GR would mainly be determined by GR’s conformation [6], GRE-driven effects are also influenced by the sequence of the GRE itself, the presence of other regulatory elements and chromatin condensation levels [6, 39, 40].

We developed three novel luciferase-based assays directly evaluating compound effects on the GR protein, namely GR dimerization (NanoBiT), protein downregulation (HiBiT-based) and Ser211 phosphorylation (BRET) (**Fig. 5**). All three assays were better predictors for compound effects on endogenous target genes than TRE-driven and GRE-driven luciferase reporters (**Fig. 6**). Quantification of GR Ser211 phosphorylation levels appeared by far the best predictor, both for direct GRE-driven activity and indirect GR-mediated downregulation of inflammatory targets. Remarkably, it was even superior to the NF-ĸB-driven reporter. The phosphorylation of Ser211 is, together with other PTMs, a very important marker for GR activity. In line with our observations (**Fig. 5c**), literature documents that the occurrence of this modification is low in absence of GR ligands, strongly induced by full agonists such as Dex and significantly less abundant in the presence of antagonists such as RU486 [31, 41]. Although reported to be important for GR’s ability to activate GRE-driven target genes [31, 32, 42, 43], this PTM also positively correlates with the pro-apoptotic actions of GR in lymphoid malignancies and with GR’s ability to downregulate mRNA levels of, among others, c-Myc, c-Jun and JunB [33, 42]. How exactly Ser211 phosphorylation leads to increased GR activity is incompletely understood, but this PTM has, together with phosphorylation of Ser203 and Ser226, been reported to result in stabilization of activation function (AF-)1 [43, 44]. This portion of the NTD is involved in the recruitment of proteins from the basal transcriptional machinery, which could help explain the importance of these particular PTMs in particular for GRE-driven signaling [44–46].

Strikingly, the ĸB-RE-driven reporter and the HiBiT-based GR-downregulation assay, both characteristic for monomer-driven GR actions [9, 12], had a higher predictive power for compound effects on the mRNA levels of GRE-driven genes than the dimer-based NanoBiT and GRE-driven reporter assays (**Fig 6a**). Inversely, NanoBiT and BRET were the best predictors for compound effects on inflammatory targets, likely subject to indirect downregulation (**Fig. 6b**). These observations were quite puzzling, especially since SEMoGRAMs and SEDiGRAMs would both be expected to change the stoichiometry of GR dimerization when compared to classic agonists, and therefore have different efficacies in assays monitoring monomer-versus dimer-driven GR activity (**Fig. 7c**). Consequently, the presence of SEMoGRAMs of SEDiGRAMs in the tested compound panel should, theoretically, have resulted in poor correlations between these two assay types. Nonetheless, this was not observed in our data. Instead, the activity profiles of the tested compounds were located on a one-dimensional plane ranging from full antagonist to full agonists, rather than being separated from one another based on how they change the stoichiometry of GR. This was confirmed by PCA, where one principal component was sufficient to capture all significant biological variation (**Fig. 7a, b**). Thus, none of these compounds appeared to be a true SEMoGRAM or SEDiGRAM, at least in A549 cells.

## Conclusion

In summary, this study reported the successful development of novel, *upstream-*positioned, GR-based assays suitable for high-throughput screening of GR ligands. Quantification of GR Ser211 phosphorylation levels proved to be a top predictor for compound activity on endogenous GR targets genes compared to the limited set of conventional luciferase reporters monitoring *downstream* GR signaling activity. In future, unbiased, multi-omics approaches should yield many additional insights, not only on how to distinguish compounds from the different functional classes, but also how the GR signaling profiles can differ between compounds from the same functional class.

## Supplementary information

Supplementary material and methods, tables and figures can be found in a separate file.

## Acknowledgements

We thank Francis Impens and Katie Boucher from the VIB Proteomics Core for the LC-MS/MS analysis. We thank Hazel Hunt from CORCEPT Therapeutics for reviewing the manuscript. Laura Van Moortel was financially supported by a strategic basic research fellowship of the Research Foundation Flanders (FWO-Vlaanderen), grant number 1S14720N.

## Abbreviations

*36B4*: acidic ribosomal phosphoprotein P0
*Angptl4*: angiopoietin-related protein 4
*AP-1*: activator protein 1
*B2M*: beta-2-microglobulin
*BRET*: bioluminescence resonance energy transfer
*CCL(2)*: chemokine (C-C motif) ligand (2)
*CNRQ*: calibrated and normalized relative quantity
*Cpd(A)*: compound (A)
*Cyclo*: cyclophilin
*DBD*: DNA-binding domain
*Dex*: dexamethasone
*DUSP1*: dual-specificity phosphatase 1
*Fam107a*: family with sequence similarity 107 member A
*FAS*: fatty acid synthase
*FKBP5*: FK506-Binding Protein 5
*Fosdagr*: fosdagrocorat
*GC*: glucocorticoid
*GILZ*: glucocorticoid-induced leucine zipper
*GR*: glucocorticoid receptor
(*n)GRE*: (negative) glucocorticoid receptor element
*GS*: glutamine synthetase
*HPRT1*: Hypoxanthine Phosphoribosyltransferase 1
*IL-1β*: interleukin 1 beta
*ĸB-RE*: NF- ĸB response element
*LBD*: ligand-binding domain
*MF*: mometasone furoate
*NF- ĸB*: nuclear factor kappa B
*NR3C1*: nuclear receptor subfamily 3 group C member 1
*NTD*: N-terminal domain
*PCA*: principal component analysis
*PEPCK*: phosphoenolpyruvate carboxykinase
*PMA*: Phorbol 12-myristate 13-acetate
*Pred*: prednisolone
*PTM*: post-translational modification
*SEGRAMs*: selective glucocorticoid receptor agonists and modulators
*SEDiGRAMs*: selective dimerizing GR agonists and modulators
*SEMoGRAMs*: selective monomerizing GR and modulators
*SGK1*: serum/glucocorticoid-regulated kinase 1
*TAT*: tyrosine aminotransferase
*TGFβ2*: transforming growth factor beta 2
*TNFα*: tumor necrosis factor alpha
*TRE*: TPA (12-O-tetradecanoylphorbol 13-acetate)-response element (= AP-1 response element)

## Statements & Declarations

### Ethical approval

Murine primary hepatocyte isolation was conducted following an approved standard protocol by the UGent Faculty of Medicine and Health Sciences Ethical Committee (ECD20-48)

### Consent to participate

Not applicable

### Consent to publish

Not applicable

### Author contributions

LVM and JT performed experiments and analyzed the data. BM and AS performed experiments. DC, DDS and SE set up experiments and interpreted data. CL and OM interpreted data. LVM, KG and KDB wrote the manuscript.

### Data availability

The mass spectrometry proteomics data have been deposited to the ProteomeXchange Consortium via the PRIDE [47] partner repository with the dataset identifier PXD031319.

