## Supplementary for "Novel assays monitoring direct glucocorticoid receptor protein activity exhibit high predictive power for ligand activity on endogenous gene targets"

### Supplementary information

*Laura Van Moortel[1,2], Jonathan Thommis[1,2], Brecht Maertens[1,2], An Staes[1,2], Dorien Clarisse[1,2], Delphine De Sutter[1,2], Claude Libert[3,4], Onno C. Meijer[5,6], Sven Eyckerman[1,2], Kris Gevaert[1,2*] & Karolien De Bosscher[1,2*]*

### Author information

**Author affiliations**

[1]VIB Center for Medical Biotechnology (CMB), Ghent, Belgium
[2]Department of Biomolecular Medicine, Ghent University, Ghent, Belgium
[3]VIB Center for Inflammation Research (IRC), Ghent, Belgium
[4]Department of Biomedical Molecular Biology, Ghent University, Ghent, Belgium
[5]Department of Medicine, Division of Endocrinology, Leiden University Medical Center, Leiden, the Netherlands.
[6]Einthoven Laboratory for Experimental Vascular Medicine, Leiden University Medical Center, Leiden, the Netherlands.

### Supplementary materials and methods

#### Indirect immunofluorescence

15,000 A549 cells were seeded in poly-L-lysine coated Bidi micro-well plates and treated for 1 h with solvent, 100 nM RU486, 10 µM CpdA or 1 µM for all other compounds. After fixation, endogenous GR was visualized using mouse anti-GR and anti-mouse AlexaFluor 488. Nuclei were stained using DAPI (300 nM, Invitrogen).

#### Immunoblot analysis

To detect Flag and GR epitopes in A549 Flag-HiBiT-GR (FH-GR) clones, cells from a confluent 12-well were lysed in 80 µL SDS sample buffer (62.5 mM Tris-HCl pH 6.8, 2% SDS, 10% glycerol, 0.01% bromophenol blue) supplemented with 50 mM dithiotreitol (DTT). To monitor GR protein levels in A549 wild-type (wt) and A549 FH-GR cells, 150,000 cells were seeded in 6-well plates and pre-incubated with solvent or 1 µM Dex in DMEM before adding mTNFα (f.c. 200 IU/mL) and incubating for 23 h. Next, cells were lysed in 200 µL SDS sample buffer supplemented with 50 mM DTT. Lysates were briefly sonicated and heated for 5 min at 95 °C before loading on a 7.5% polyacrylamide gel. After SDS-PAGE, proteins were transferred to 0.45-µm nitrocellulose membrane (GE Healthcare) for 2 h at 100V or overnight at 30 V (4 °C). Membranes were blocked using StartingBlock TBS blocking buffer (Thermo Scientific) mixed 1:1 with TBS-T (50 mM Tris-HCl pH 7.5, 150 mM NaCl supplemented with 0.1% Tween20) and subsequently incubated with antibodies to Flag (1:2000), GR (1:2000) and β-actin (1:30,000). After washing steps in TBS-T, membranes were incubated with HRP-linked whole antibodies (GE Healthcare) before detection on an Amersham Imager 680 using the Vector® NovaRED peroxidase (HRP) substrate kit (Vector Laboratories).

#### Sample preparation for LC-MS/MS

1.5 x $10^6$ A549 cells were seeded in 10-cm plates, and pre-treated with compounds for 1 h before adding mTNFα (2,000 IU/mL) for 5 h. Cells were collected by scraping in DPBS (with $Ca^{2+}$ and $Mg^{2+}$) and cell pellets were snap-frozen in liquid $N_2$ and stored at -80 °C until further use. Pellets were lysed in 100 µL 10% SDS and 100 mM TEAB (pH 8.5), sonicated for 3 times for 10 s with 30 s on ice in between and centrifuged for 15 min at 20,000 g. Protein concentrations were determined using Pierce™ BCA Protein Assay (ThermoFisher) and samples were diluted to 2 mg/mL. 100 µg protein material (50 µL) was consecutively incubated with 15 mM DTT (30 min at 55 °C), 30 mM iodoacetamide (IAA, 15 min at room temperature (RT) in the dark) and acidified to pH 1 by adding 10 µL 12% phosphoric acid. Next, 350 µL 90% methanol (MeOH) in 100 mM triethylammonium bicarbonate (TEAB) pH 7.55 was added, samples were transferred to an S-trap plate and centrifuged for 2min at 1,500 g. The plate was washed 3 times with 200 µL 90% methanol in 100 mM TEAB pH 7.55, followed by a 2-min centrifugation at 1,500 g, and proteins were subsequently digested overnight at 37 °C by adding 1 µg trypsin in 125 µL 50 mM TEAB. The next day, peptides were consecutively washed with 8 µL 50 mM TEAB, 80 µL 0.2% formic acid and 80 µL 50% acetonitrile in 0.2% formic acid, each time followed by a 2-min centrifugation at 1,500 g. Finally, peptides were transferred to a mass spectrometry vial, dried in a SpeedVac™ vacuum concentrator and stored at -20 °C until further use. Right before injection, each sample was solubilized in 20 µL loading solvent A (0.1% TFA in water:acetonitrile (ACN) (98:2, v:v)).

#### LC/MS-MS and data analysis

2 µg peptide material was injected for LC-MS/MS analysis on an Ultimate 3000 RSLCnano system in-line connected to an Orbitrap Fusion Lumos mass spectrometer (Thermo). Trapping was performed at 10 µL/min for 4 min in loading solvent A on a 20 mm trapping column (made in-house, 100 µm internal diameter (I.D.), 5 µm beads, C18 Reprosil-HD, Dr. Maisch, Germany). The peptides were separated on a 50 cm µPAC™ Gen2 column (C18-endcapped functionality, 300 µm wide channels, 2.5 µm porous-shell pillars, inter pillar distance of 1.25 µm and a depth of 3 µm; PharmaFluidics, Belgium). It was kept at a constant temperature of 50 °C. Peptides were eluted at a flow rate of 250 nL/min by a linear gradient starting from solvent A, reaching 5% solvent B (0.1% formic acid (FA) in water/acetonitrile (2:8, v/v)) after 10 min, 22.5% solvent B after 109 min and 30.5% solvent B at 135 min, 55% solvent B at 153 min, 70% solvent B at 155 min, followed by a 5-minutes wash at 70% MS solvent B and re-equilibration with MS solvent A (0.1% FA in water). The mass spectrometer was operated in data-dependent mode, automatically switching between MS and MS/MS acquisition in TopSpeed mode with a cycle time of 3s. Full-scan MS spectra (300-1500 m/z) were acquired at a resolution of 120,000 in the Orbitrap analyzer after accumulation to a target AGC value

of 200,000 with a maximum injection time of 250ms. The precursor ions were filtered for charge states (2-7 required), dynamic exclusion (60 s; ± 10 ppm window) and intensity (minimal intensity of 5E3). The precursor ions were selected in the ion routing multipole with an isolation window of 1.2 Da and accumulated to an AGC target of 12E3 or a maximum injection time of 40 ms and activated using HCD fragmentation (34% normalized collision energy (NCE)). The fragments were analyzed in the Ion Trap Analyzer at normal scan rate. The polysiloxane peak at m/z 445.12003 was used as an internal lock mass.

Data searching was done with the MaxQuant software (v1.6.11.0) using the Andromeda search engine with default search settings including a false-discovery rate (FDR) of 1% on both the peptide and protein level. Spectra were searched against the human SwissProt proteome database (version of January 2021). The mass tolerance for precursor and fragment ions was set to 20 ppm and 4.5 ppm, respectively, during the main search. Enzyme specificity was set as C-terminal to arginine and lysine (trypsin), also when followed by a Pro residue, with a maximum of 2 missed cleavages. Variable modifications were set to oxidation of methionine residues and acetylation of the protein N-terminus. A minimum of one razor or unique peptide was required for identification. Matching between runs was enabled with a 20-min alignment window and a matching time window of 0.7 min. Proteins were quantified by the MaxLFQ algorithm integrated in the software, with the fastLFQ switched off and a minimum ratio count of 2 unique or razor peptides. Further data analysis was performed with the Perseus software (v.1.6.2.1) using the ProteinGroups table from the MaxQuant search output. Potential contaminants and proteins identified in a reverse database were removed, as well as proteins that were only identified by site. The protein LFQ intensities were log2-transformed to obtain a normal distribution, and proteins with less than 70% valid values across all samples were removed. The biological replicates were grouped by their treatment. Two sample t-tests were performed versus solvent condition, with a permutation-based FDR of 0.05 and S0 of 0.1 for truncation. The results were visualized as volcano plots.

### Supplementary data

**Supplementary Table 1** overview compounds

| *Name* | *Structure* | *Described ligand class* | *Predicted ligand class (based on* **Fig. 1***)* | *References* |
|---|---|---|---|---|
| | | *Controls* | | |
| CompoundA (CpdA) | 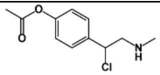 | SEGRAM | GR-independent | [1–6] |
| Dexamethasone (Dex) | 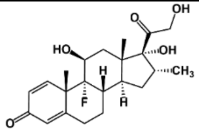 | Full agonist | Full agonist | [7–9] |
| | | *Test compounds* | | |
| AL-438 | 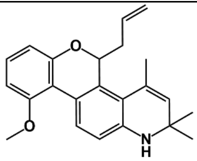 | SEGRAM | Partial agonist | [10–14] |
| AZD2906 | 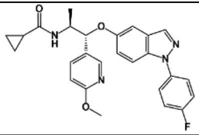 | SEGRAM | SEDiGRAM | [15–18] |
| Compound1 (Cpd1) | 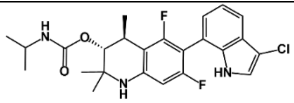 | SEGRAM | Full agonist | [19] |
| Compound2 (Cpd2) | 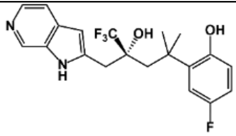 | SEGRAM | Partial agonist | [20–23] |
| Compound3 (Cpd3) | 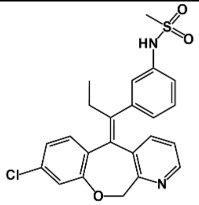 | SEGRAM | SEMoGRAM | Structural analogues hereof in [24] |
| Compound4 (Cpd4) | undisclosed | undisclosed | SEDiGRAM | / |
| Compound5 (Cpd5) | 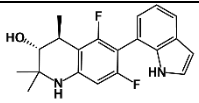 | Partial agonist | SEDiGRAM | [19, 25] |
| CORT108297 | 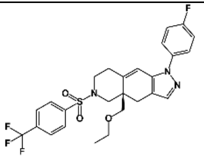 | SEGRAM | SEMoGRAM | [26–33] |
| CORT118335 | 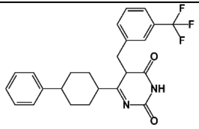 | SEGRAM | Antagonist | [26, 27, 33–37] |

| | | | | |
|---|---|---|---|---|
| Cortisol | 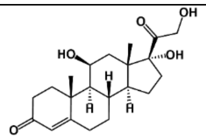 | Full agonist | Full agonist | [7, 38, 39] |
| Cortivazol | 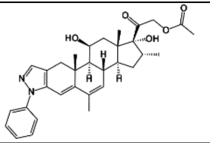 | Full agonist | Full agonist | [15, 40–46] |
| Dagrocorat<br>PF-(00251)802 | 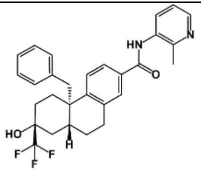 | SEGRAM | SEMoGRAM | [47–49] |
| Fosdagrocorat<br>(Fosdagr)<br>PF-04171327 | 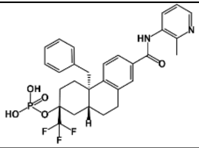 | SEGRAM | SEMoGRAM | [15, 47, 50–55] |
| LGD5552 | 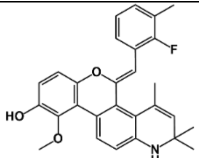 | Partial agonist/SEGRAM | Partial agonist | [15, 56, 57] |
| Mapracorat | 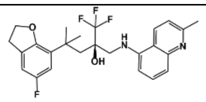 | SEGRAM | Full agonist | [15, 58–66] |
| MK-5932 | 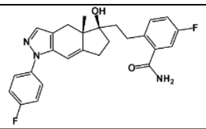 | SEGRAM | SEMoGRAM | [67, 68] |
| Momethasone<br>Furoate<br>(MF) | 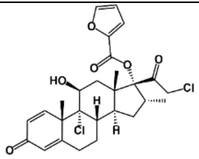 | Full agonist | Full agonist | [69–71] |
| PF-(04308)515 | 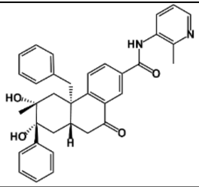 | SEGRAM | SEMoGRAM | [47] |
| Prednisolone<br>(Pred) | 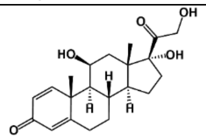 | Full agonist | Full agonist | [15] |
| RU486 | 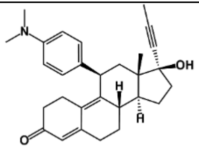 | Antagonist | Antagonist | [72, 73] |

**Supplementary Table 2** Primer list

| | | |
|---|---|---|
| | *CRISPR-Cas9 knock-in* | |
| | *Forward* | *Reverse* |
| Guide RNA | CACCGACCAGGAGTTAATGATTCTT | AAACAAGAATCATTAACTCCTGGTC |
| PCR of NR3C1 terminus | GTGAGCGGCAGGATGAATAG | CCCGAGGAAAGGCTGATTTG |
| Ultramer single stranded DNA | TTATGATGTTTTCCCCCCGTTTTTGTTTTTTGTTTTGTAGTTGATATTCACTGATGGACTACAAAGACGATGACGACAAGGCGATGGTGAGCGGCTGGCGGCTGTTCAAGAAGATTAGCGCTAGTGACTCTAAAGAATCATTAACTCCTGGTAGAGAAGAAAACCCCAGCAGTGTGCTTGCTCAGGAGAGG | |
| | *Human qPCR primers* | |
| *Target* | *Forward* | *Reverse* |
| *h36B4* | CATGCTCAACATCTCCCCCTTCTCC | GGGAAGGTGTAATCCGTCTCCACAG |
| *hAngptl4* | GGCTCAGTGGACTTCAACCG | CCGTGATGCTATGCACCTTCT |
| *hCCL2* | CAGCCAGATGCAATCAATGCC | TGGAATCCTGAACCCACTTCT |
| *hCCL5* | TGCCCACATCAAGGAGTATTT | TTTCGGGTGACAAAGACGA |
| *hCyclo* | ATGGTGATCTTCTTGCTGGTCCTTGC | GCATACGGGTCCTGGCATCTTGTCC |
| *hDUSP1* | CTGCCTTGATCAACGTCTCA | CTGTGCCTTGTGGTTGTCCT |
| *hFKBP5* | AATGGTGAGGAAACGCCGATG | TCGAGGGAATTTTAGGGAGACT |
| *hGILZ* | GCGTGAGAACACCCTGTTGA | TCAGACAGGACTGGAACTTCTCC |
| *hGR* | TGATGAAGCTTCAGGATGTCA | TTCGAGCTTCCAGGTTCATTC |
| *hGS* | GGGGACTAATGCCGAGGTC | ACGATGCAAGATGAAACGGG |
| *hHPRT1* | TGACACTGGCAAAACAATGCA | GGTCCTTTTCACCAGCAAGCT |
| *hIL-1β* | GAGCTACGAATCTCCGACCAC | GGCAGGGAACCAGCATCTTC |
| *hIL-8* | GCTCTCTTGGCAGCCTTCCTGA | ACAATAATTTCTGTGTTGGCGC |
| *hSGK1* | GAGATTGTGTTAGCTCCAAAGC | CTGTGATCAGGCATACCACACT |
| *hTGFβ2* | AGAGTGCCTGAACAACGGATT | CCATTCGCCTTCTGCTCTT |
| *hTNFα* | | |
| | *Murine qPCR primers* | |
| *Target* | *Forward* | *Reverse* |
| *mAngptl4* | GGAAAGAGGCTTCCCAAGATG | CGTTGGGAGTCAAGCCAATG |
| *mB2M* | CATGGCTCGCTCGGTGAC | CAGTTCAGTATGTTCGGCTTCC |
| *mDUSP1* | AGTACCCCTCTCTACGATCAGG | CGAGAAGCGTGATAGGCACTG |
| *mFam107a* | TCATCAAACCCAAGAAGCTG | CTCAGGCTTGCTGTCCATAC |
| *mFAS* | GGTATGTCCGGGAAATTGCC | ATTGTGTGTGCCTGCTTGGG |
| *mFKBP5* | TGAGGGCACCAGTAACAATGG | CAACATCCCTTTGTAGTGGACAT |
| *mGILZ* | GGCCCTAGACAACAAGATTGAG | CACGAATCTGCTCCTTTAGGAC |
| *mGR* | TGGAGAGGACAACCTGACTTCC | ACGGAGGAGAACTCACATCTGG |
| *mHPRT1* | CCTAAGATGAGCGCAAGTTGAA | CCACAGGACTAGAACACCTGCTAA |
| *mPEPCK* | TGACAGACTCGCCCTATGTG | CCCAGTTGTTGACCAAAGGC |
| *mSGK1* | GAGATCGTGTTAGCTCCAAAGC | CTGTGATCAGGCATAGCACACT |
| *mTAT* | TGCTGGATGTTCGCGTCAATA | CGCCTTCACCTTCATGTTGTC |

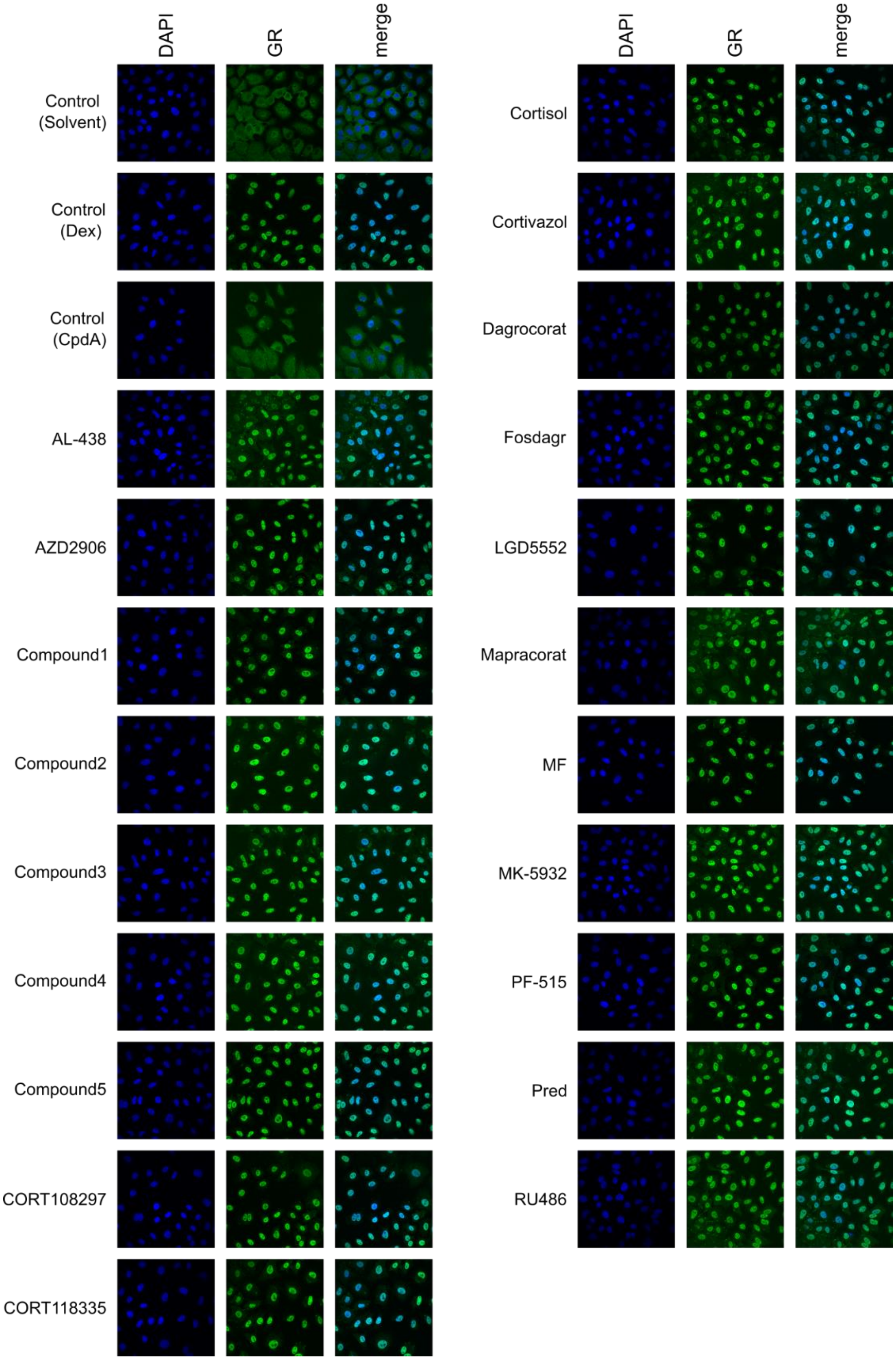


**Supplementary Fig. 1** Evaluation of GR nuclear translocation via indirect immunofluorescence. A549 cells were treated with compounds for 1h. Endogenous GR was visualized using anti-GR and AlexaFluor488 (green), nuclei were stained with DAPI (blue).

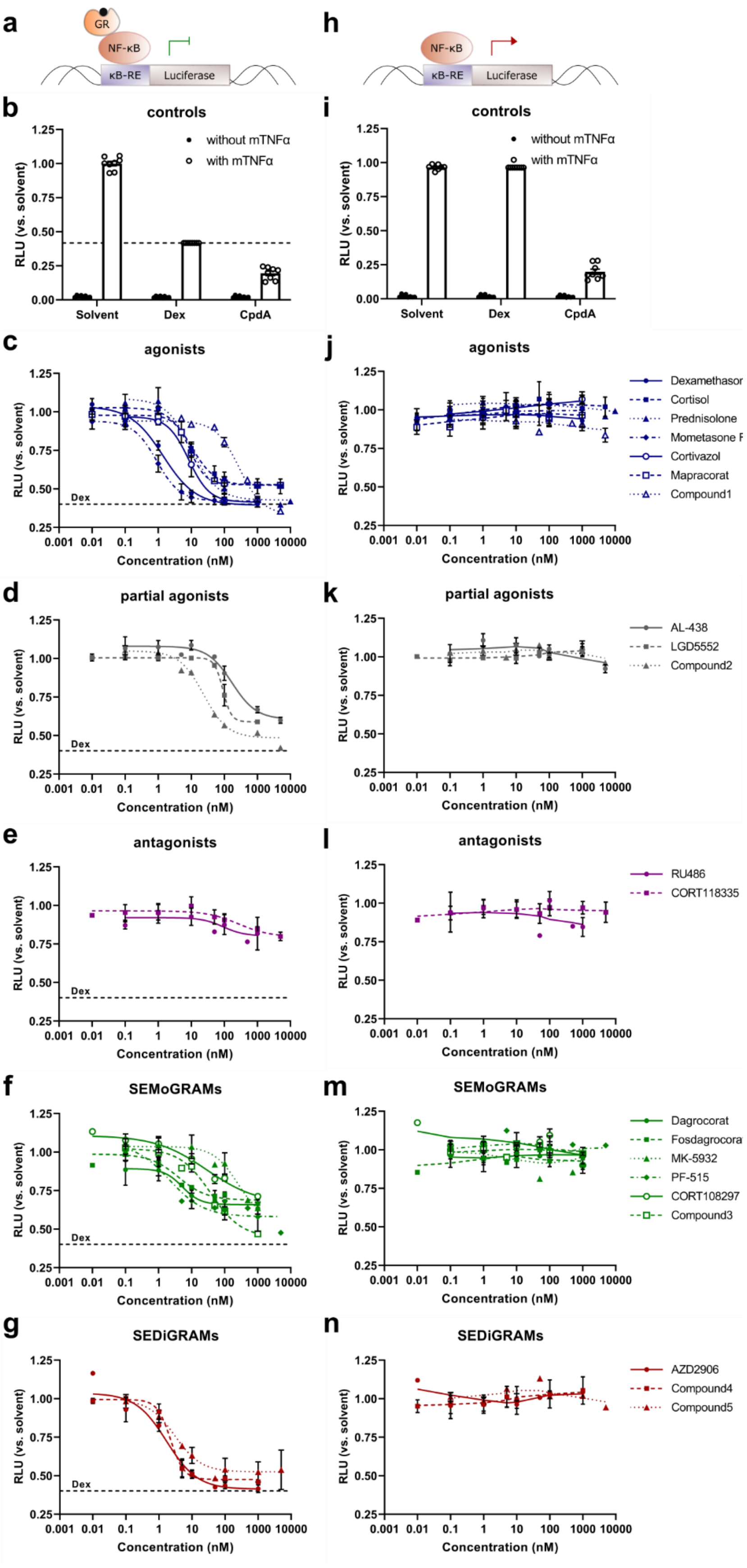
a
GR
NF-κB
κB-RE
Luciferase
h
NF-κB
κB-RE
Luciferase
b
controls
without mTNFα
with mTNFα
RLU (vs. solvent)
Solvent
Dex
CpdA
i
controls
without mTNFα
with mTNFα
RLU (vs. solvent)
Solvent
Dex
CpdA
c
agonists
Dex
RLU (vs. solvent)
Concentration (nM)
j
agonists
Dexamethasor
Cortisol
Prednisolone
Mometasone F
Cortivazol
Mapracorat
Compound1
d
partial agonists
Dex
RLU (vs. solvent)
Concentration (nM)
k
partial agonists
AL-438
LGD5552
Compound2
e
antagonists
Dex
RLU (vs. solvent)
Concentration (nM)
l
antagonists
RU486
CORT118335
f
SEMoGRAMs
Dex
RLU (vs. solvent)
Concentration (nM)
m
SEMoGRAMs
Dagrocorat
Fosdagrocora
MK-5932
PF-515
CORT108297
Compound3
g
SEDiGRAMs
Dex
RLU (vs. solvent)
Concentration (nM)
n
SEDiGRAMs
AZD2906
Compound4
Compound5

**Supplementary Fig. 2** GR-dependency of compound-induced NF-κB suppression. Wild-type (**a-g**) or GR knock-out (**h-n**) A549 cells, stably transfected with $(NF\text{-}\kappa B)_3$_Luc, were seeded in 96-well plates and treated with compound serial dilutions, solvent, 1 µM Dex or 10 µM CpdA for 6 h. After 1h, cells were induced with mTNFα (2000 IU/mL). All conditions were tested in technical replicates and data were normalized versus 1 µM Dex for each plate separately. (**a-g**) The results from at least 3 biological replicates were used for nonlinear curve fittings in GraphPad Prism 9 (4 parameters). (**h-n**) the results from at least 3 biological replicates were subjected to smoothing in GraphPad Prism 9 (4 neighbors on each side, second order)

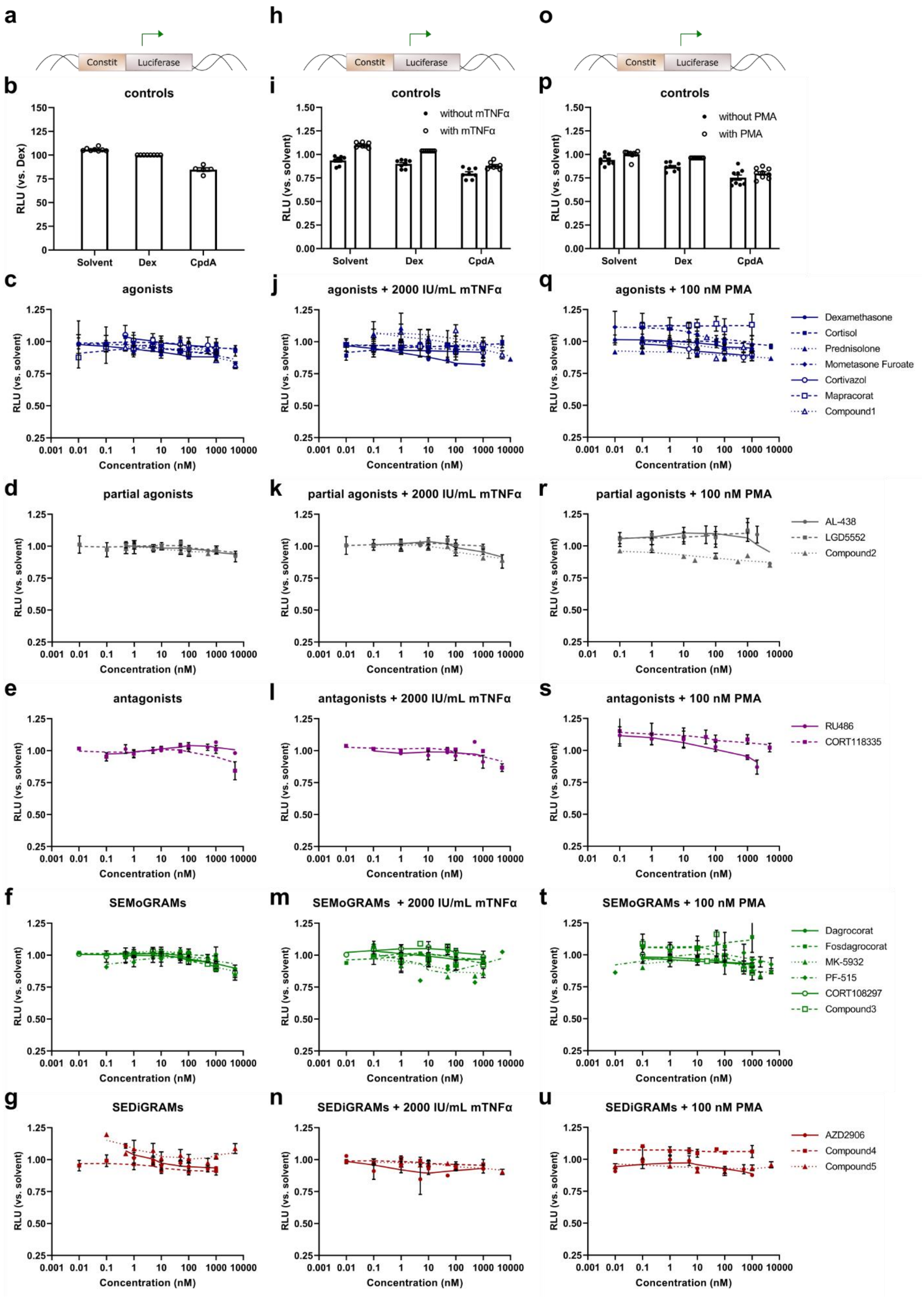
a
h
o
Constit
Luciferase
b
controls
RLU (vs. Dex)
Solvent
Dex
CpdA
i
controls
without mTNFα
with mTNFα
RLU (vs. solvent)
p
controls
without PMA
with PMA
c
agonists
j
agonists + 2000 IU/mL mTNFα
q
agonists + 100 nM PMA
Dexamethasone
Cortisol
Prednisolone
Mometasone Furoate
Cortivazol
Mapracorat
Compound1
Concentration (nM)
d
partial agonists
k
partial agonists + 2000 IU/mL mTNFα
r
partial agonists + 100 nM PMA
AL-438
LGD5552
Compound2
e
antagonists
l
antagonists + 2000 IU/mL mTNFα
s
antagonists + 100 nM PMA
RU486
CORT118335
f
SEMoGRAMs
m
SEMoGRAMs + 2000 IU/mL mTNFα
t
SEMoGRAMs + 100 nM PMA
Dagrocorat
Fosdagrocorat
MK-5932
PF-515
CORT108297
Compound3
g
SEDiGRAMs
n
SEDiGRAMs + 2000 IU/mL mTNFα
u
SEDiGRAMs + 100 nM PMA
AZD2906
Compound4
Compound5

**Supplementary Fig. 3** Monitoring of compound effects on cell viability. A549 cells, stably transfected with constitutively expressed luciferase (Const_Luc) were seeded in 96-well plates and treated with compound serial dilutions, solvent, 1 µM Dex or 10 µM CpdA for 6 h. After 1h, cells were induced with mTNFα (2000 IU/mL) (**h-n**) or 100 nM PMA (**o-u**). All conditions were tested in technical replicates and data were normalized versus 1 µM Dex for each plate separately. The results from at least 3 biological replicates were subjected to smoothing in GraphPad Prism 9 (4 neighbors on each side, second order)

**Supplementary Table 3** Overview of calculated compound efficacies ($E_{max}$), potencies ($EC_{50}$ or $IC_{50}$) and R squared from nonlinear curve fittings on the data represented in **Fig. 2**. Absence of the 95% confidence interval (CI) marked as ?? indicates that this could not be calculated. * no best-fit value could be found, efficacy of the highest non-toxic compound concentration

| | *A549 GRE_Luc* | | | *A549 (NF-κB)₃_Luc* | | | *A549 Collagenase3_Luc* | | | |
|---|---|---|---|---|---|---|---|---|---|---|
| | *$E_{max}$ (% vs. Dex) [95% CI]* | *$EC_{50}$ [95% CI]* | *$R^2$* | *$E_{max}$ (% vs. Dex) [95% CI]* | *$IC_{50}$ [95% CI]* | *$R^2$* | *$E_{max}$ (% vs. Dex) [95% CI]* | *$IC_{50}$ [95% CI]* | *$R^2$* | *$E_{max}$ GRE/ $E_{max}$ κB-RE* |
| *Dex* | 100 [97.3 - 102.9] | 6.71 [5.85 - 7.61] | 1.00 | 100 [92.9 - 109.5] | 1.4 [0.74 - 2.35] | 0.97 | 100 [85.1 - 140.7] | 2.58 [0.63 - 8.91] | 0.90 | 1.00 |
| *Cortisol* | 77.1 [71.9 - 82.5] | 34.28 [25.66 - 44.56] | 0.98 | 78.6 [70.1 - 88.8] | 9.95 [5.73 - 17.57] | 0.95 | 95.0 [76.4 - ??] | 32.66 [10.42 - 272.3] | 0.82 | 0.98 |
| *Pred* | 87.1 [82.8 - 91.8] | 34.37 [26.84 - 43.03] | 0.98 | 94.5 [81.1 - 114.0] | 10.34 [4.79 - 24.96] | 0.93 | 101.0 [91.1 - 112.1] | 19.86 [?? - 40.13] | 0.94 | 0.92 |
| *MF* | 106.8 [99.9 - 113.9] | 2.26 [1.37 - 3.18] | 0.96 | 95.5 [87.2 -105.1] | 0.90 [0.43 - 1.59] | 0.94 | 94.2 [83.5 - 105.5] | 1.88 [?? to 3.02] | 0.91 | 1.12 |
| *Cortivazol* | 92.0 [81.6 - 104.4] | 19.89 [11.77 - 34.53] | 0.92 | 96.7 [85.2 - 114.1] | 8.3 [3.32 - 16.10] | 0.94 | 98.5 [87.2 - 110.0] | 4.29 [2.50 - 6.25] | 0.92 | 0.95 |
| *Cpd1* | 84.5 | ~ 502.6 | 0.96 | 109.9 [99.3 - 123.8] | 280.5 [193.8 - 413.1] | 0.98 | 99.1* | 299.3 [68.69 - ??] | 0.94 | 0.77 |
| *AL-438* | 36.1 [31.7 - 41.6] | 207.1 [131.5 - 327.3] | 0.92 | 65.2 [50.0 - 114.7] | 170.6 [87.55 - 991.2] | 0.90 | 74.7* | N.D. | 0.87 | 0.55 |
| *LGD5552* | 44.4 [40.5 - 53.0] | 127.3 [?? - 361.0] | 0.95 | 67.9 [58.1 - 77.7] | 90.6 [72.20 - 115.8] | 0.93 | 88.7 [74.6 - 112.1] | 61.1 [35.85 - 105.1] | 0.92 | 0.65 |
| *Cpd2* | 51.2 [46.1 - 56.4] | 45.45 [30.68 - 57.34] | 0.94 | 85.1 [78.2 - 92.9] | 23.44 [17.06 - 32.61] | 0.98 | 96.1 [81.3 - ???] | 53.37 [22.49 - ??] | 0.89 | 0.60 |
| *RU486* | 5.3 [4.7 - 6.0] | 41.44 [17.84 - 73.20] | 0.74 | 32.7 | 92.06 | 0.18 | 79.7* | N.D. | 0.88 | 0.16 |
| *CORT118335* | 8.3 [4.7 - ??] | N.D. | 0.11 | 33.7 | 267.9 | 0.35 | 41.7 | N.D. | 0.62 | 0.25 |
| *Dagrocorat* | 17.4 | 48.03 [23.28 - 54.07] | 0.93 | 84.4 [45.7 - ??] | 5.189 | 0.71 | 49.5 | 4.821 | 0.82 | 0.31 |
| *Fosdagr* | 17.2 | N.D | 0.93 | 51.5 [43.5 - 73.7] | 3.98 [1.26 - 16.74] | 0.81 | 67.2* | N.D. | 0.59 | 0.33 |
| *Mapracorat* | 95.6 [88.8 - 104.1] | 12.34 [?? - 18.31] | 0.97 | 78.0 [69.2 - 89.3] | 9.21 [5.92 - 16.97] | 0.91 | 71.1 [47.4- 131.0] | 45.94 [?? - 280.4] | 0.73 | 1.23 |
| *MK-5932* | 12.8 [11.9 - 13.9] | 297.6 [218.8 - ??] | 0.97 | 73.7 | 250.2 [55.93 - ??] | 0.77 | 65.9 [46.5 - ??] | 251 [95.44 - ??] | 0.77 | 0.17 |
| *PF-515* | 15.9 [13.8 - 19.9] | 6.30 [0.83 - 14.78] | 0.88 | 69.3 [55.2 - ??] | 2.14 | 0.79 | 61.6* | N.D. | 0.57 | 0.23 |
| *CORT108297* | 24.2 [20.4 - 35.7] | 99.57 [47.65 - 585.1] | 0.78 | 55.8 [37.7 - ??] | 28.57 [5.83 - ??] | 0.86 | 51.2* | N.D. | 0.36 | 0.43 |

| | | | | | | | | | | |
|---|---|---|---|---|---|---|---|---|---|---|
| *Cpd3* | 41.0 [35.1 - 47.1] | 59.84 [43.48 - 79.48] | 0.89 | 91.5 [67.0 - 155.0] | 37.62 [15.62 - 201.5] | 0.92 | 108.5 [83.1 - ??] | 62.36 [26.42 - ??] | 0.87 | 0.45 |
| *AZD2906* | 134.7 [126.0 - 145.3] | 12.5 [9.88 - 18.38] | 0.97 | 97.0 [85.6 - 126.3] | 1.70 [0.023 - 3.59] | 0.90 | 100.2 [87.2 - 113.4] | 4.62 [2.31 - 6.95] | 0.88 | 1.39 |
| *Cpd4* | 135.4 [119.5 - 151.9] | 3.76 [2.25 - 5.77] | 0.93 | 86.7 [80.8 - 92.8] | 2.24 [?? - 3.18] | 0.97 | 79.5 [64.3 - 95.1] | 3.01 [?? - 5.03] | 0.74 | 1.56 |
| *Cpd5* | 106.2 [99.9 - 112.6] | 13.81 [10.31 - 18.54] | 0.98 | 78.5 [66.6 - 101.3] | 3.04 [0.00004 - 9.68] | 0.76 | 113.4 [100.3 - 147.1] | 12.44 [5.275 - 49.97] | 0.95 | 1.35 |

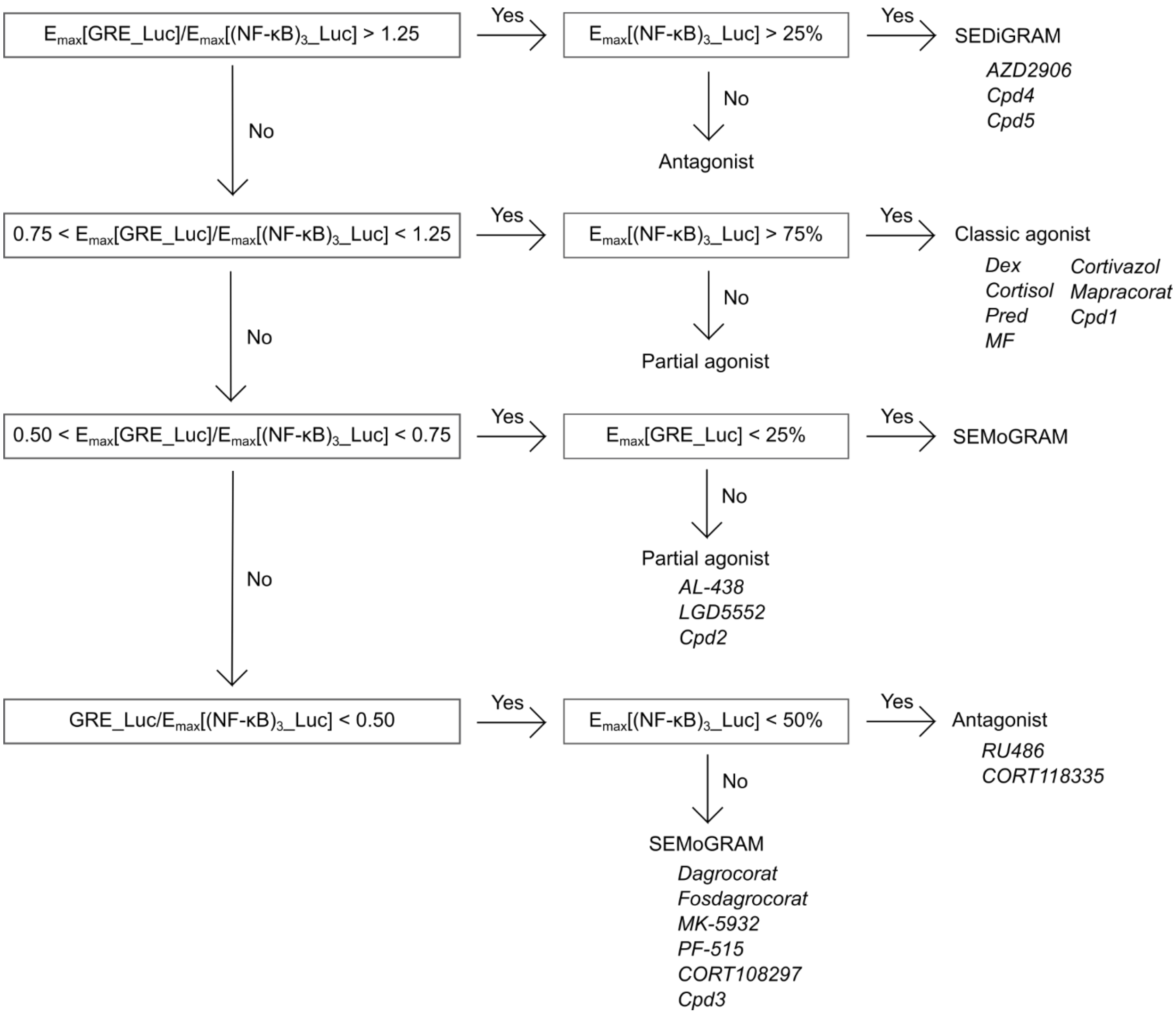


**Supplementary Fig. 4** Flowchart for compound classification based on their efficacies ($E_{max}$) on $(NF\text{-}\kappa B)_3$_Luc and GRE_Luc (**Fig. 2, Supplementary Table 3**).

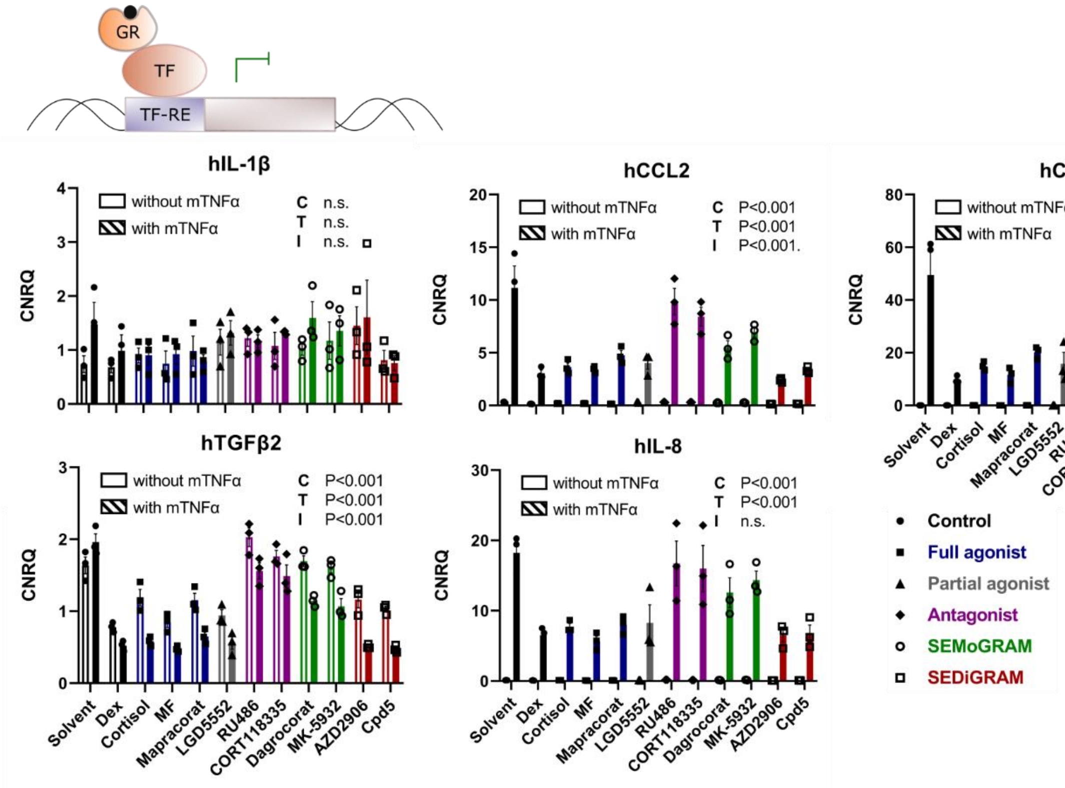


**Supplementary Fig. 5** Compound effects on mRNA levels of pro-inflammatory makers in A549 in absence and presence of mTNFα. A549 cells were pre-incubated with compounds for 1 h, before adding mTNFα (2000 IU/mL) for 5 h. Total mRNA was isolated, reverse transcribed and used for qPCR analysis. Graphs represent the mRNA levels inflammatory markers subject to indirect GR-mediated downregulation (3 biological replicates), normalized versus hHPRT1, hCyclo and h36B4 mRNA levels. Indirect GR-mediated downregulation of inflammatory markers. Significant changes versus solvent and Dex were evaluated via 2-way ANOVA followed by Dunnett's post-hoc testing and are reported in **Supplementary Table 4 and 5.** C, compound effect in 2-way ANOVA; T, mTNFα effect in 2-way ANOVA; I, 2-way ANOVA interaction factor

**Supplementary Table 4** Overview of statistically significant differences between solvent and compounds on the mRNA levels of GR target genes in A549 cells (**Fig. 3**, **Supplementary Fig. 5**). Significance levels were calculated via Dunnett's multiple comparison tests, following 2-way ANOVA. *, P < 0.05; **, P < 0.01; ***, P < 0.001; ns, not significant; n.a., not applicable

| | *hAngptl4* | *hFKBP5* | *hGS* | *hSGK1* | *hDUSP1* | *hGILZ* | *hTNFAIP3* | *hGR* | *hIL-1β* | *hCCL2* | *hCCL5* | *TGFβ2* | *hIL-8* | *hTNFα* |
|---|---|---|---|---|---|---|---|---|---|---|---|---|---|---|
| *without mTNFα* | | | | | | | | | | | | | | |
| Dex | *** | *** | *** | *** | *** | *** | *** | ns | ns | *** | ns | *** | ns | n.a. |
| Cortisol | *** | *** | *** | *** | *** | *** | *** | ns | ns | ** | ns | * | ns | n.a. |
| MF | *** | *** | *** | *** | *** | *** | *** | * | ns | ** | ns | *** | * | n.a. |
| Mapracorat | *** | *** | * | *** | *** | *** | *** | ns | ns | *** | ns | * | ns | n.a. |
| LGD5552 | *** | *** | ns | * | *** | *** | *** | ns | ns | ns | ns | *** | ns | n.a. |
| RU486 | *** | *** | ns | ns | *** | * | ** | ns | ns | ns | ns | ns | ns | n.a. |
| CORT118335 | *** | *** | ns | ns | *** | ** | ** | ns | ns | ns | ns | ns | ns | n.a. |
| Dagrocorat | *** | *** | ns | ns | *** | *** | ** | ns | ns | ns | ns | ns | ns | n.a. |
| MK-5932 | *** | *** | ns | * | *** | *** | *** | ns | ns | ns | ns | ns | ns | n.a. |
| AZD2906 | *** | *** | *** | *** | *** | *** | *** | ns | ns | *** | ns | * | ns | n.a. |
| Cpd5 | *** | *** | *** | *** | *** | *** | *** | ns | ns | *** | ns | ** | ns | n.a. |
| *with mTNFα* | | | | | | | | | | | | | | |
| Dex | *** | *** | *** | *** | *** | *** | * | ns | ns | *** | *** | *** | *** | *** |
| Cortisol | *** | *** | *** | *** | *** | *** | ns | ns | ns | *** | ** | *** | ** | *** |
| MF | *** | *** | *** | *** | *** | *** | * | ns | ns | *** | *** | *** | *** | *** |
| Mapracorat | *** | *** | ns | *** | *** | *** | ns | ns | ns | *** | ns | *** | ** | ** |
| LGD5552 | *** | *** | ns | ns | *** | *** | ns | * | ns | *** | ** | *** | ** | *** |
| RU486 | *** | ns | ns | ns | * | ns | ns | ns | ns | ns | ns | ns | ns | ns |
| CORT118335 | *** | *** | ns | ns | *** | ns | ns | ns | ns | ns | ns | ns | ns | ns |
| Dagrocorat | *** | *** | ns | ns | *** | *** | ns | ns | ns | *** | ns | *** | ns | * |
| MK-5932 | *** | *** | ns | ns | *** | *** | ns | ns | ns | * | ns | *** | ns | ns |
| AZD2906 | *** | *** | *** | *** | *** | *** | ns | * | ns | *** | *** | *** | *** | *** |
| Cpd5 | *** | *** | *** | *** | *** | *** | ns | ns | ns | *** | ** | *** | *** | *** |

**Supplementary Table 5** Overview of statistically significant differences between Dex and compounds on the mRNA levels of GR target genes in A549 cells (**Fig. 3**, **Supplementary Fig. 5**). Significance levels were calculated via Dunnett's multiple comparison tests, following 2-way ANOVA. *, P < 0.05; **, P < 0.01; ***, P < 0.001; ns, not significant; n.a., not applicable

| | *hAngptl4* | *hFKBP5* | *hGS* | *hSGK1* | *hDUSP1* | *hGILZ* | *hTNFAIP3* | *hGR* | *hIL-1β* | *hCCL2* | *hCCL5* | *TGFβ2* | *hIL-8* | *hTNFα* |
|---|---|---|---|---|---|---|---|---|---|---|---|---|---|---|
| *without mTNFα* | | | | | | | | | | | | | | |
| Solvent | *** | *** | *** | *** | *** | *** | *** | ns | ns | *** | ns | *** | ns | n.a. |
| Cortisol | ns | ns | ns | ns | ns | ns | ns | ns | ns | ns | ns | ** | ns | n.a. |
| MF | ns | ns | ns | ns | ns | ns | ns | ns | ns | ns | ns | ns | ns | n.a. |
| Mapracorat | ns | ns | ns | ns | ns | ns | ns | ns | ns | ns | ns | ** | ns | n.a. |
| LGD5552 | ns | ns | ** | *** | ns | ns | ns | ns | ns | *** | ns | ns | ns | n.a. |
| RU486 | *** | *** | *** | *** | *** | *** | ** | ns | ns | *** | ns | *** | * | n.a. |
| CORT118335 | *** | *** | *** | *** | *** | *** | ** | * | ns | *** | ns | *** | ns | n.a. |
| Dagrocorat | *** | ** | * | *** | * | *** | ** | ns | ns | ** | ns | *** | ns | n.a. |
| MK-5932 | *** | * | ns | *** | ns | *** | ns | ns | ns | ** | ns | *** | ns | n.a. |
| AZD2906 | ns | ns | * | ns | ns | ns | ns | ns | ns | ns | ns | ** | ns | n.a. |
| Cpd5 | ns | ns | ns | ns | ns | ns | ns | ns | ns | ns | ns | ns | ns | n.a. |
| *with mTNFα* | | | | | | | | | | | | | | |
| Solvent | *** | *** | *** | *** | *** | *** | * | ns | ns | *** | *** | *** | *** | *** |
| Cortisol | ns | ns | ns | ns | ns | ns | ns | ns | ns | ns | ns | ns | ns | ns |
| MF | ns | ns | ns | ns | ns | ns | ns | ns | ns | ns | ns | ns | ns | ns |
| Mapracorat | ** | ns | *** | *** | ns | * | ns | ns | ns | * | ns | ns | ns | ns |
| LGD5552 | *** | *** | *** | *** | *** | * | ** | ns | ns | ns | ns | ns | ns | ns |
| RU486 | *** | *** | *** | *** | *** | *** | ns | ns | ns | *** | *** | *** | ** | *** |
| CORT118335 | *** | *** | *** | *** | *** | *** | ns | ** | ns | *** | *** | *** | ** | ** |
| Dagrocorat | *** | *** | *** | *** | *** | *** | ns | ns | ns | ** | ns | *** | * | ns |
| MK-5932 | *** | *** | *** | *** | ** | *** | ns | ns | ns | *** | ** | *** | ** | * |
| AZD2906 | ns | ns | ns | ns | ns | ns | ns | ns | ns | ns | ns | ns | ns | ns |
| Cpd5 | ns | ns | ns | ns | ns | ns | ns | ns | ns | ns | ns | ns | ns | ns |

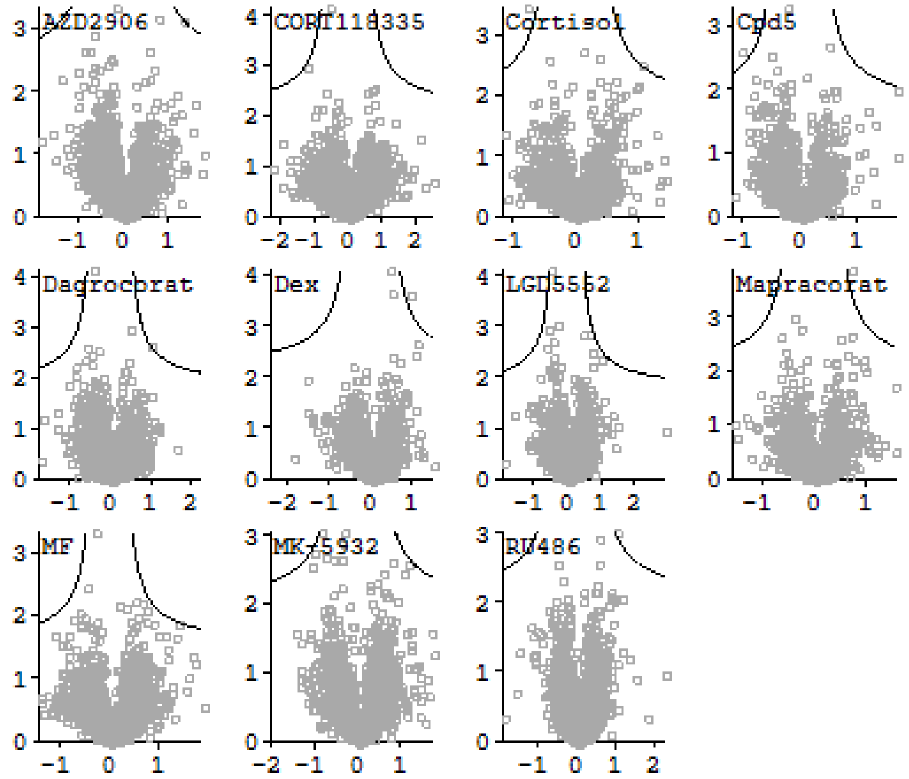


**Supplementary Fig. 6** Evaluation of compound effects on the proteome of A549 cells. A549 cells were treated with compounds for 1 h before adding mTNFα (2000 IU/mL) for 5 h. Cells were collected and lysed, proteins were digested with trypsin and the resulting peptides were analysis via LC-MS/MS. Volcano plots represent compound effects versus solvent. 4331 proteins were identified, common contaminants, only identified by site and reverse identifications not included. Compound effects versus solvent were evaluated via multiple two-sample t-tests with a permutation-based FDR of 0.05 and S0 value of 0.1.

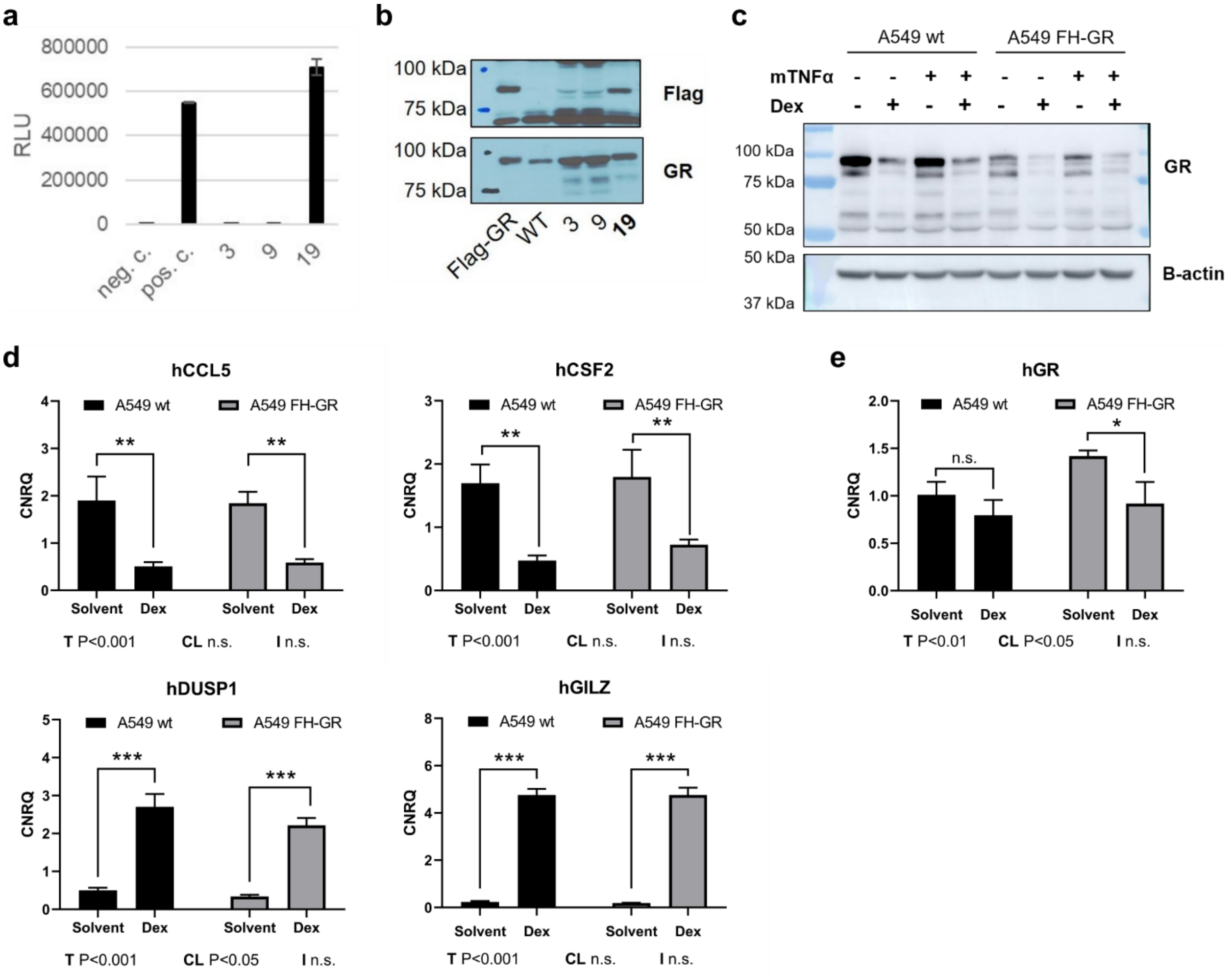


**Supplementary Fig. 7** Validation of A549 Flag-HiBiT-GR knock-in cell line clone 19 (A549 FH-GR). **(a)** Confirmation of the presence of functional HiBiT tag via measurement of NanoLuc signal. **(b)** Detection of Flag-tag and endogenous GR via immunoblot analysis. **(c)** Monitoring of protein GR levels in A549 wild-type (wt) and A549 FH-GR cells, evaluated via Western blotting. Cells were incubated with solvent or 1 µM Dex for 1 h before adding mTNFα (200 IU/mL) for 23 h. **(d)** mRNA levels of GR target genes in A549 wt versus A549 FH-GR cells (3 biological replicates), normalized versus hHPRT1 and hCyclo. Cells were incubated with solvent or 1 µM Dex for 1 h before adding mTNFα (2000 IU/mL) for 5 h. **(e)** hGR mRNA levels in A549 wt versus A549 FH-GR cells (3 biological replicates), normalized versus h36B4 and hCyclo. Cells were incubated with solvent or 1 µM Dex for 1 h before adding mTNFα (200 IU/mL) for 23 h. **(d, e)** Significant changes were evaluated via 2-way ANOVA followed by Tukey's multiple comparisons test. T, treatment effect in 2-way ANOVA; CI, cell line effect in 2-way ANOVA; I, 2-way ANOVA interaction factor

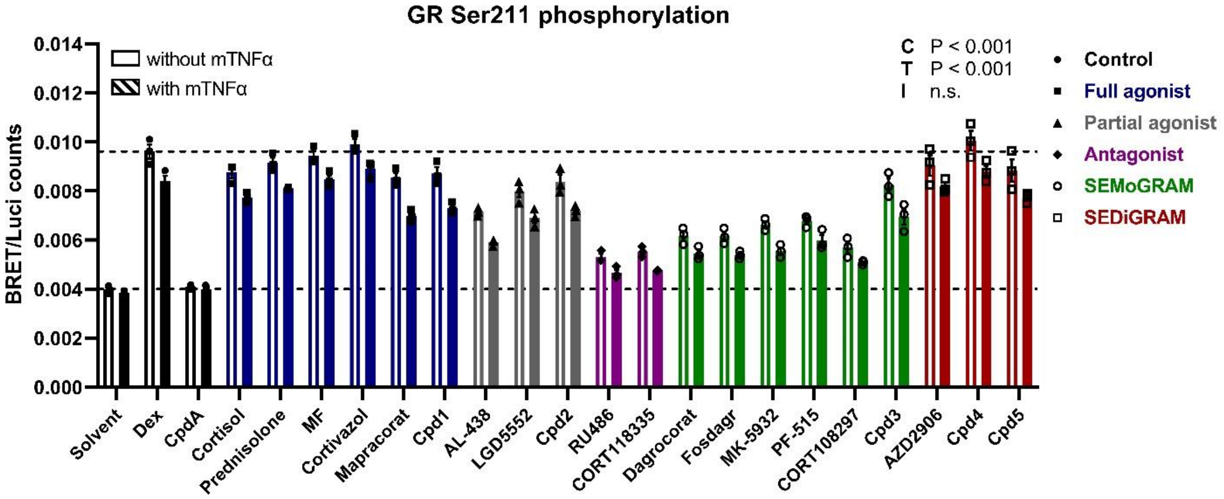


**Supplementary Fig. 8** Quantification of GR Ser211 phosphorylation levels via BRET. A549 Flag-HiBiT-GR cells were pre-incubated with compounds for 1h, followed by 1h mTNFα (2000 IU/mL) induction where indicated. Lysates were incubated with anti-GR and LgBiT for 30 min, followed by addition of furimazine substrate and anti-rabbit Alexa Fluor 594 for 60 min. BRET/NanoLuc ratios were calculated for each well separately and used for further analysis. All conditions were measured in technical triplicates. Statistically significant changes versus Dex were evaluated via 2-way ANOVA with Dunnett's post-hoc testing on log2-transformed data (3 biological replicates). Results from post-hoc testing are provided in **Supplementary Table 6.** RLU, relative luciferase units; C, compound effect in 2-way ANOVA; T, mTNFα effect in 2-way ANOVA; I, 2-way ANOVA interaction factor

**Supplementary Table 6** Statistical analysis of compound effects on GR Ser211 phosphorylation in A549 FH-GR (**Fig. 5c**). Significance levels were calculated via 2-way ANOVA followed by Dunnett's multiple comparisons tests versus solvent and Dex conditions. *, P < 0.05; **, P < 0.01; ***, P < 0.001; ns, not significant

| | | | | |
|---|---|---|---|---|
| *2-way ANOVA* | | | | |
| Compound | *** | | | |
| mTNFα | *** | | | |
| Interaction factor | ns | | | |
| *Multiple comparisons* | | | | |
| | *Versus solvent* | | *Versus Dex* | |
| | *No mTNFα* | *With mTNFα* | *No mTNFα* | *Without mTNFα* |
| Solvent | / | / | *** | *** |
| Dex | *** | *** | / | / |
| CpdA | ns | ns | *** | *** |
| Cortisol | *** | *** | ns | ns |
| Prednisolone | *** | *** | ns | ns |
| MF | *** | *** | ns | ns |
| Cortivazol | *** | *** | ns | ns |
| Mapracorat | *** | *** | * | *** |
| Cpd1 | *** | *** | ns | ** |
| AL-438 | *** | *** | *** | *** |
| LGD5552 | *** | *** | *** | *** |
| Cpd2 | *** | *** | ** | ** |
| RU486 | *** | *** | *** | *** |
| CORT118335 | *** | *** | *** | *** |
| Dagrocorat | *** | *** | *** | *** |
| Fosdagr | *** | *** | *** | *** |
| MK-5932 | *** | *** | *** | *** |
| PF-515 | *** | *** | *** | *** |
| CORT108297 | *** | *** | *** | *** |
| Cpd3 | *** | *** | ** | *** |
| AZD2906 | *** | *** | ns | ns |
| Cpd4 | *** | *** | ns | ns |
| Cpd5 | *** | *** | ns | ns |

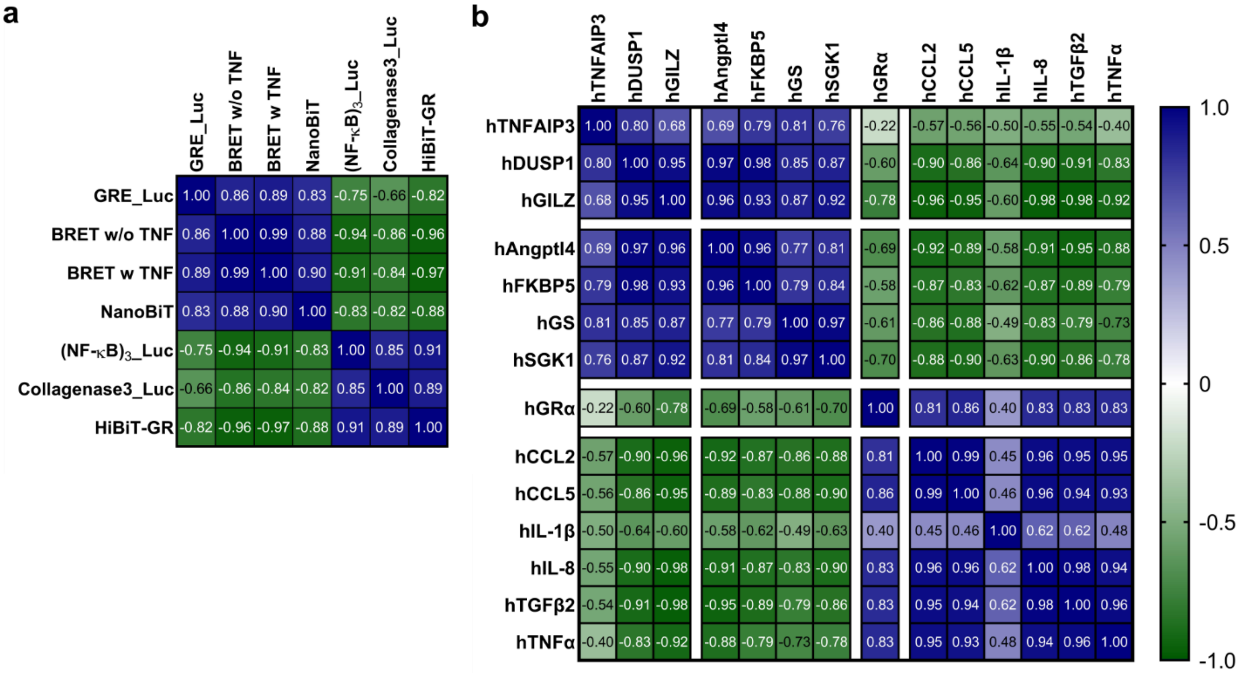


**Supplementary Fig. 9** Correlation studies of the data reported in **Fig. 2-5**. (**a**) Heatmap representing the Pearson correlations between the luciferase-based assays on all 20 compounds (**b**) Heath map representing the Pearson correlations between all tested qPCR target genes. (**a, b**) Pearson correlations were calculated based on the mean values of the log2-transformed data in GraphPad Prism 9.
